# Multimodal spatial transcriptomics determines repeat expansion, huntingtin aggregation, and selective cortical neuron loss in Huntington’s Disease

**DOI:** 10.64898/2025.12.26.696622

**Authors:** Sonia Vazquez-Sanchez, Olatz Arnold-Garcia, Carlos Chillon-Marinas, Roy Maimon, Katelyn McNair, Yuanyi Dai, Natalia Jimenez-Villegas, Jess Cui, Leslie M. Thompson, Jone Lopez-Erauskin, Bogdan Bintu, Don W. Cleveland

**Affiliations:** Department of Cellular and Molecular Medicine, University of California at San Diego, La Jolla, CA, USA; Department of Neurosciences, Biogipuzkoa Health Research Institute, San Sebastián 20014, Spain; Centro de Investigación Biomédica en Red sobre Enfermedades Neurodegenerativas (CIBERNED), Carlos III Institute (ISCIII), Spanish Ministry of Sciences and Innovation, Madrid 28029, Spain; Department of Biomedical Engineering, New York University, Tandon School of Engineering, NY, NY, USA; Department of Neurobiology & Behavior, University of California, Irvine, Irvine, CA 92617, USA; Institute for Memory Impairments and Neurological Disorders, University of California, Irvine, Irvine, CA 92617, USA; Department of Biological Chemistry, University of California, Irvine, Irvine, CA 92617, USA; Sue and Bill Gross Stem Cell Research Center, University of California, Irvine, Irvine, CA 92617, USA; Center for Neurotherapeutics, University of California, Irvine, Irvine, CA 92617, USA; Department of Neurology, University of California, Irvine, Irvine, CA 92617, USA; Department of Psychiatry & Human Behavior, University of California, Irvine, Irvine, CA 92617, USA; Department of Neurosciences, University of California at San Diego, La Jolla, CA, USA

## Abstract

Huntington’s disease (HD) is caused by CAG expansion in *HTT*, yet how somatic repeat instability and huntingtin aggregation relate to selective cell loss in the human brain remains unclear. We have developed a multimodal spatial transcriptomics approach that enables defining transcriptional programs with subcellular resolution, somatic CAG repeat lengths, and six other pathology marks including huntingtin aggregates in every cell of intact brain sections. Imaging 428,173 cells in HD cortex revealed selective vulnerability: L5–6 NP and L6b deep-layer excitatory neurons undergo >50% loss, closely linked to very large (>380±55) somatic expansions. Intranuclear aggregation was most prevalent at intermediate somatic repeat expansion (220-300 CAGs) and was accompanied by broader transcriptional changes. In contrast, chandelier and somatostatin+ inhibitory interneurons are lost despite only modest repeat expansion or aggregation. These data provide a comprehensive resource and establish a broadly applicable framework for connecting repeat expansion and protein pathology across diverse cell types.

**Short Bullet points:**

- Development of a novel, multimodal spatial transcriptomics platform enables definition of RNA transcriptomes with subcellular resolution, somatic repeat expansions, and protein accumulation in cells within tissue sections
- Generation of two complementary datasets for HD and control cortex: 428,173 cells to quantify comprehensively cell-type vulnerability, and a deeper multimodal dataset in which CAG repeat expansion, huntingtin aggregation, six additional pathological readouts, and expression of 1,128 genes were measured in 185,721 cells, providing a comprehensive resource for the neurodegeneration community.
- Deep layer excitatory neuron loss (L5–6 NP and L6b) was associated with very large somatic CAG expansions (>380±55), while selective inhibitory neuron loss (chandelier and somatostatin interneurons) occurred with modest CAG repeat expansion or huntingtin intranuclear aggregation
- Intranuclear aggregation, not somatic repeat expansion, was more predictive of transcriptional changes, including chromatin remodeling and RNA export factors

## INTRODUCTION

Huntington’s Disease (HD) is a neurodegenerative disorder caused by a CAG expansion in the huntingtin gene (*HTT*), with more than 40 CAG repeats in exon 1 of *HTT* being fully penetrant^1,2^. The expanded CAG tract encodes an elongated polyglutamine stretch that promotes mutant huntingtin misfolding and aggregation^1,2^. Human HD brains show abundant nuclear and cytoplasmic inclusions composed of huntingtin protein fragments^3,4^. Mutant huntingtin protein disrupts multiple cellular processes—including RNA splicing^5–7^ and transport^8,9^. Recent large-scale profiling of HD patients has revealed that genomic CAG repeat length, rather than polyglutamine length, is the strongest predictor of disease onset^10,11^. Furthermore, many modifier genes identified in genome-wide association studies (GWAS) are involved in DNA mismatch repair and influence somatic repeat instability, supporting a model in which toxicity arises, at least in part, from ongoing CAG expansion in vulnerable cells^10,12,13^. While both HTT protein aggregation and somatic repeat expansion appear to contribute to pathogenesis^1,14^, measuring their molecular manifestation at single-cell resolution and how they relate to cell vulnerability in the human brain remains challenging.

Anatomically, HD is characterized by prominent degeneration in the striatum and cortex^1,2^. Striatal degeneration underlies motor symptoms while cortical degeneration contributes to cognitive decline and psychiatric disturbances ^1,2^. At the molecular level, however, it remains unclear what are the most affected cell types and within them what transcriptional programs change in relation to HTT aggregation and somatic repeat expansion. Single-nuclear RNA sequencing coupled with long-read sequencing of *HTT* has revealed that large somatic repeat expansions occur in medium spiny neurons (MSNs)—the striatal cells that die most prominently—and are associated with transcriptional dysregulation^15^. This work focused exclusively on the striatum and the approach introduced^15^, along with other related similar methods^16,17^, are powerful but, are limited in bridging the anatomical and molecular scales. Given the limited spatial and molecular resolution, these approaches lack the systematic profiling of all neuronal and non-neuronal subtypes to identify which cells in intact human brain sections are selectively lost, define their transcriptional alterations, and their relationship to the two leading pathogenic mechanisms—huntingtin aggregation and somatic repeat expansion.

New multiplexed spatial transcriptomics imaging methods, especially Multiplex Error-Robust Fluorescence in-situ Hybridization (MERFISH)^18–20^, have been developed to permit determining RNA levels of hundreds to thousands of selected genes within individual cells^21,22^. Unlike single-cell (or single nuclei) RNA sequencing methods in which cells or nuclei are first isolated thereby losing spatial resolution, the MERFISH approach retains spatial context and morphological information of the cells within an intact tissue section. This is achieved through error-robust barcoding, combinatorial labeling (using ∼50 encoding probes per RNA), and sequential rounds of single-molecule FISH imaging—typically completed in 10–20 imaging cycles^19^. Here, we report development of an extension to the prior MERFISH approaches^23,24^, referred here as multimodal MERFISH, which permits determination of CAG repeat expansions, expression levels of transcriptional programs at subcellular resolution, and evaluation of protein localization and expression within every cell in an intact tissue section. We applied this approach to the human HD and control cortex, profiling 1,128 genes alongside somatic CAG repeat expansion in *HTT* and the accumulation of huntingtin and six additional proteins. This integrated dataset allowed us to determine the cells most vulnerable in HD cortex and directly relate intranuclear huntingtin aggregation, repeat expansion, transcriptional remodeling, and DNA repair gene expression at single-cell resolution.

## RESULTS

### Cell profiling of the HD frontal cortex using Multimodal Spatial Transcriptomics

To address how CAG somatic repeat expansion and HTT aggregation impact single-cells in the human brain, we developed a multimodal spatial transcriptomics approach (called here Multimodal MERFISH) that enables comprehensive profiling of cell type, determination of transcriptional programs, detection of subcellular protein localization (including huntingtin (HTT) and HTT associated cytoplasmic or nuclear aggregates), and somatic CAG repeat expansions in *HTT* RNA within each cell of an intact tissue section (**Figure 1A-C and S1**). We performed Multimodal MERFISH to map 1128 genes (**MM-1128,Table S1**) in frontal cortex sections from five HD donors and four donors with non-neurological disease used as controls (**Table S2**). The library includes 357 cell-type markers (based on the Allen’s Institute’s public snRNAseq dataset^25^), genes covering 222 markers of inflammation or neuronal death, 59 annotated genes of the DNA repair machinery and 558 members of the pathways affected in HD, including 66 for RNA splicing^5^ and 225 for nuclear transport^8,9^. Each encoding probe was synthesized with an acrydite modification to enable covalent incorporation into a thin polyacrylamide gel that was cast directly onto the tissue sample, ensuring long-term probe retention. Each of our imaging experiments began and ended with imaging of the single transcripts of two marker genes - *NRGN* and *GAD1* by smFISH – confirming a 94.7% signal retention after a total of 72 hybridization and imaging rounds covering all the 1128 genes targeted (**Figure S1F**).

**Figure 1:**
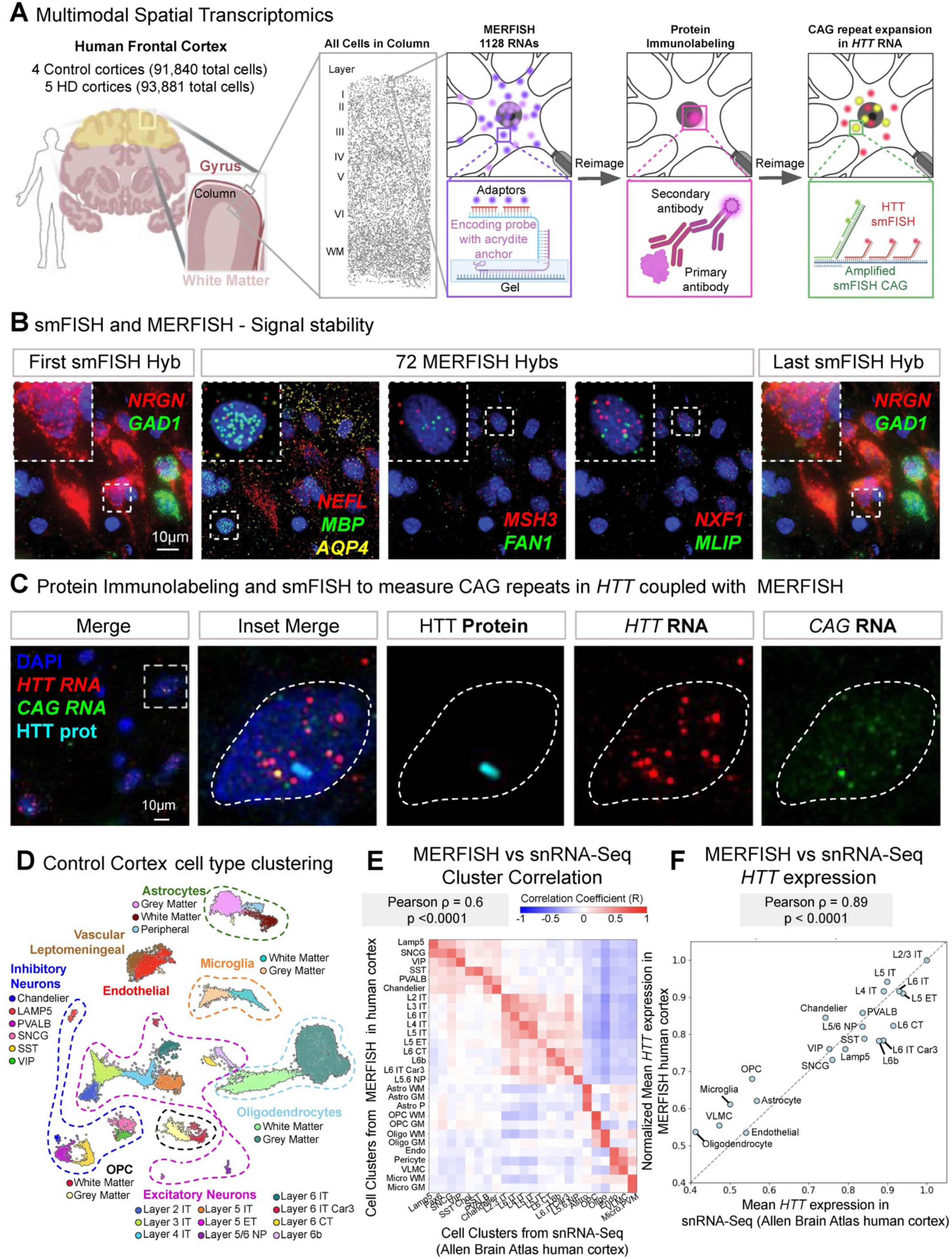
Multimodal MERFISH measures transcriptomes, somatic CAG repeat expansion in Huntingtin RNAs, and accumulation of pathology readouts including aggregates within every cell of an intact section of HD brain. (A) Multimodal MERIFSH overview, a platform which measures the degree of *HTT* CAG expansion at single cell resolution in postmortem cortical sections, along with transcriptomes from 1128 genes and the intracellular localization of 8 target proteins including HTT. (B) Representative images of smFISH for *NRGN* (red) in the first (left) and last (right) cycle of hybridization and imaging. Representative images of MERFISH decoded transcripts imaged across 72 cycles of hybridization and imaging within the same field of view. DAPI in blue. (C) Protein immunolabeling and smFISH to measure CAG repeats in *HTT* coupled with MERFISH. (D) UMAP representation of 29,456 cells and 1128 genes in human control cortex. (E) Heatmap Pearson correlation plots comparing the major cell type clusters in our MERFISH data (6 non-neurological control and 8 HD frontal cortices) and snRNA-seq performed by the Allen Brain Atlas. (F) Correlation between mean HTT expression in each cell cluster in our Multimodal MERFISH data compared with data from Allen’s Institute’s public snRNAseq dataset^25^.

Cell segmentation was performed using machine learning methods^26^ as previously described^19,21^ (**Methods**) and then cell types were determined based on single-cell transcriptional measurements using the Leiden clustering algorithm^27^. An example from one non-neurological control cortical sample covering 29,456 cells, allowed for defining 28 cell type clusters (**Figure 1D**). Our MERFISH-defined cell clusters were consistent with prior single-nuclear sequencing atlases of the human cortex^25^ (ρ = 0.66, p = <0.0001). **(Figure 1E** and **S2-S3)** and with established molecular cell type markers^16^ **(Figure S2)**. Expression levels of specific genes, including *HTT,* across cell types were also consistent with those reported by single-nuclear sequencing^25^ (ρ = 0.89, p = <0.0001, **Figure 1F**). Remarkably, while the cell clusters were defined exclusively by their single-cell RNA expression profiles measured by MERFISH, there were multiple examples where each cluster was restricted to a specific cortical spatial location (**Figure S4-S5**). For example, excitatory neurons of different classes were confined to their corresponding cortical layers while oligodendrocytes separated into two subclusters: one in the white matter, and another in the gray matter (**Figure S4-S5**).

To quantify cell type abundance across the HD cortex, we adapted stereology methods from histopathology^28^ to our molecularly defined spatial maps (see Methods). Briefly, we segmented the cortex into equally wide columns spanning from the surface of the cortex to the white matter and quantified the number of cell types within these regions (**Figure 2A** and **Supplementary Figures S6A**). To extend our quantification of cell-type vulnerability to a larger cohort, we imaged four additional HD and four additional control cortical samples, using a reduced Multimodal MERFISH library (**MM-188, Table S1**) covering 188 genes selected from the larger library (MM-1128) for accurate cell type definition. While the larger library (MM-1128) covered more transcriptional programs, the cell types defined by the two probe sets matched the single-nuclear sequencing results nearly identically (ρ = 0.66, p = <0.0001 **Figure 1E** and ρ = 0.65, p = <0.0001 **Figure S3**).

**Figure 2:**
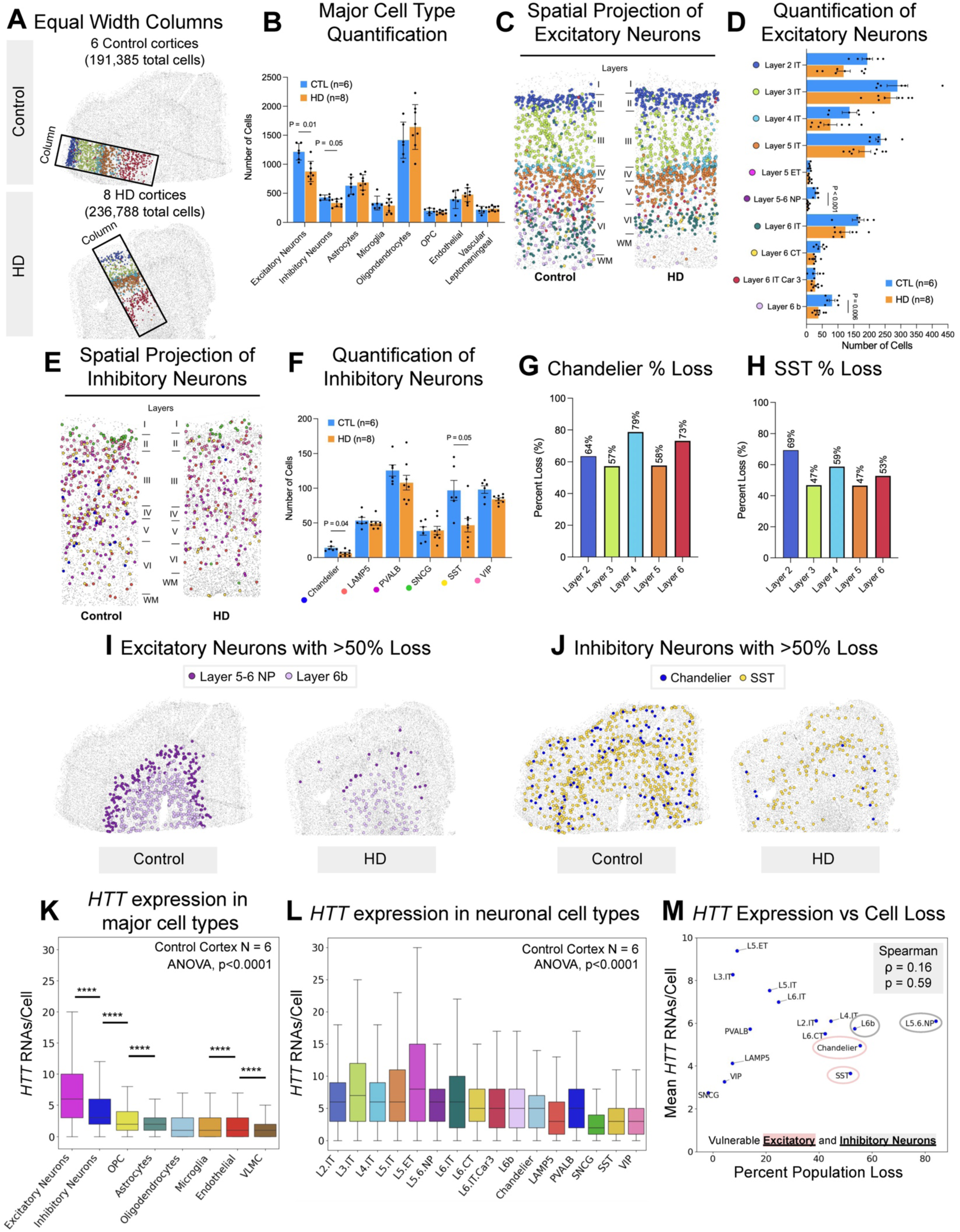
Multimodal MERFISH reveals more than 50% neuronal loss in excitatory subpopulations of neurons of layers 5 and 6 (L5-6 NP and L6b) and within Chandelier and somatostatin positive inhibitory neurons across all layers. **(A)**. Equal width columns with height defined between Layer 1/peripheral astrocytes to the white matter astrocytes are used to quantify cell numbers between control and HD cortices. (**B**) Number of cells in the mayor cell type clusters in controls versus HD. (**C**) Spatial map of non-neurological control and HD frontal cortices showing the excitatory neurons subtypes. (**D**) Quantification of the number of excitatory neurons subtypes in 6 non-neurological control and 8 HD frontal cortices. (**E**) Spatial map of non-neurological control and HD frontal cortices showing the inhibitory neurons subtypes from (C) with matching colors. (**F**) Quantification of the number of inhibitory neurons subtypes in 6 non-neurological control and 8 HD frontal cortices. (**G**) Quantification of the percentage loss of Chandelier inhibitory neurons across cortical layers. (**H**) Quantification of the percentage loss of SST-positive inhibitory neurons across cortical layers. (**I**) Spatial map of non-neurological control and HD frontal cortices showing the excitatory neuron subtypes most decrease in HD cortex (Layer 5-6NP and layer 6b) with matching colors. (**J**) Spatial map of non-neurological control and HD frontal cortices showing the inhibitory neuron subtypes most decrease in HD cortex (Chandelier and SST) with matching colors. (**K**) Total HTT transcript per cell in the major cell type clusters in control cortices. (**L**) Total HTT transcript per cell in the excitatory neuron subtype clusters in control cortices. (**M**) Total HTT transcript per cell in the major cell type clusters in control and HD cortices.

The cell abundance quantification revealed that both inhibitory and excitatory neurons were reduced, albeit overall modestly, in the HD cortex compared to controls (p = 0.01 and p = 0.05, respectively) **(Figure 2B)**. Non-neuronal cell populations remained largely unchanged. Interestingly microglia showed no signs of proliferation **(Figure 2B** and **Supplementary Figures S6F-G)**, in contrast to the robust microgliosis reported in Alzheimer’s disease^29^. Despite the modest overall changes in inhibitory and excitatory neurons at the broader class level, we observed high variability in neuronal loss across the different neuronal subtypes. Layer 5–6 Near-Projecting (L5–6 NP) neurons were the most affected subtype, showing an 84% reduction (p < 0.0001), consistent with recent reports^16^, followed by Layer 6b neurons, which exhibited a 53% reduction (p = 0.006) **(Figure 2C-D, I** and **S6D)**. This differential neuronal loss across different excitatory neuron subtypes was reproducible across cortical samples and was more pronounced in individuals with advanced disease stage (higher Vonsattel grade^30^) (**Figure S6B**). Subclasses of inhibitory neurons were also significantly reduced in HD cortex, with chandelier cells and somatostatin-expressing (*SST*) interneurons each showing approximately 50% reductions throughout all cortical layers (p = 0.04 and p = 0.05, respectively) (**Figure 2E-H, J, S6E** and **S7**), particularly in advanced disease stages (**Figure S6C**).

We observed differential *HTT* expression across the different neuronal subpopulations (**Figure 2K**), potentially reflecting differences in their physiological demand for *HTT*. To investigate whether *HTT* expression levels is predictive of the differential vulnerability observed across the different neuronal subtypes, we quantified *HTT* transcript levels per cell in control cortices. Although expression levels varied across neuronal subtypes (**Figure 2L**), *HTT* transcript abundance per cell type did not correlate with the degree of neuronal loss observed in HD cortex (ρ =0.16, p = 0.59) (**Figure 2M**), suggesting that *HTT* expression is not a primary determinant of selective vulnerability.

### Intranuclear HTT aggregates are not enriched in the neuronal types with highest cell loss

We developed a protocol permitting the combination of MERFISH with immunolabeling of HTT protein (**Figure 3A**) using two antibodies whose epitopes lie in the N-terminus prior to or and after the polyglutamine track, respectively (**Figure S8A**). This approach allowed us to quantify both intranuclear and extranuclear HTT accumulations, previously referred to as “aggregates”^31^ (**Figure 3A-B**). In HD cortex, aggregates were detected both within nuclei (intranuclear aggregates: ∼15%) and in the cytoplasm or extracellular space (extranuclear aggregates: ∼85%) (**Figure S8B-C**). Extranuclear aggregates were more abundant and significantly larger in volume (extranuclear: 50 µm³, radius ≈ 2.29 µm; intranuclear: 40 µm³, radius ≈ 2.08 µm; p < 0.0001) (**Figure S8D**) and enriched in the deeper cortical layers (**Figure 3C-F** and **S8E-F**).

**Figure 3:**
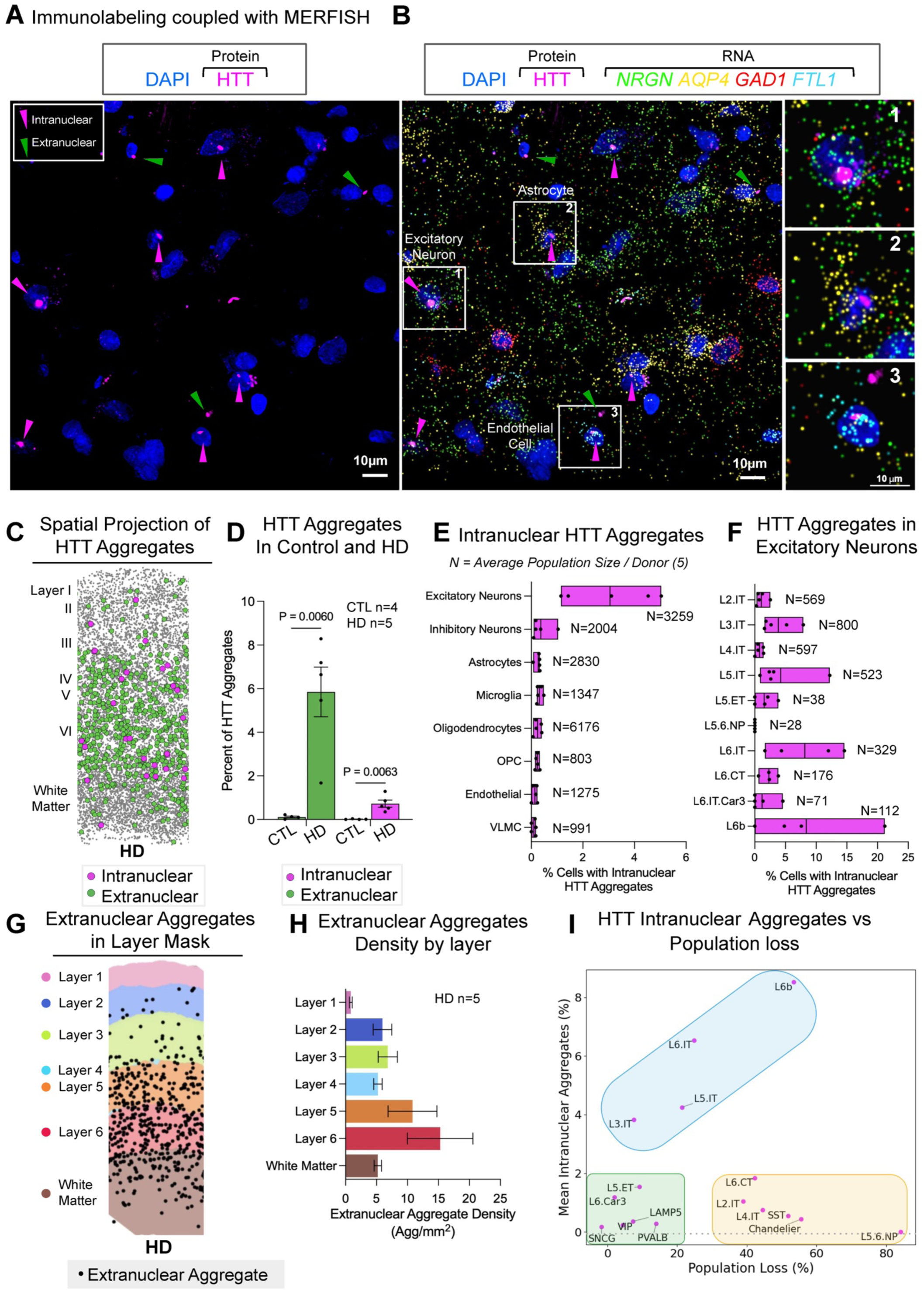
Intranuclear HTT aggregates are not enriched in the cell type most affected by cell loss. (**A**) Representative images of immunofluorescent labelling of huntingtin protein (magenta). Magenta arrowheads indicate intranuclear huntingtin aggregates and green arrowheads extranuclear. DAPI in blue. (**B**) Same field of view as in (A) aligned with MERFISH decoded transcripts for excitatory neuron gene *NRGN* (green), astrocytic maker *AQP4* (yellow), inhibitory neuron marker *GAD1* (red) and endothelial marker *FTL1* (cyan). (**B**) Spatial map of HD frontal cortex depicting intranuclear (magenta) and extranuclear (green) huntingtin aggregates. (**C**) Spatial map of HD frontal cortex depicting extranuclear aggregates (black) in different cortical layers. (**D**) Percentage of cells with intranuclear and extranuclear HTT aggregates in control and HD cortex. (**E**) Spatial map of HD frontal cortex depicting extranuclear aggregate density. (**F**) Extranuclear aggregate density in each cortical layer. (**G**) Percentage of cells with HTT intranuclear aggregates. (**G**) Percentage of excitatory neurons subtypes with HTT intranuclear aggregates. (**H**) Extranuclear HTT extranuclear aggregate density in the different cortical layers. (**I**) Correlation between the percentage of neurons with intranuclear aggregates in each cell cluster in our Multimodal MERFISH data compared with percentage of cells lost in HD cortex.

While intranuclear aggregates were less frequent than extranuclear ones **(Figure 3D** and **S8B-C)**, mutant HTT within the nucleus has been suggested to disrupt transcription, RNA processing, and DNA integrity^2,5,9^. Using automated fluorescence intensity-based detection of HTT inclusions, combined with 3D nuclear segmentation from DAPI signal—validated by Lamin A and NUP98 co-staining (**Figure S8G-H**)—we found that intranuclear aggregates were predominantly localized to excitatory neurons, a pattern consistently observed across all five HD cortical samples analyzed (**Figure 3G**). Interestingly, Layer 5–6 Near-Projecting (L5–6 NP) neurons—despite being the most vulnerable to cell loss—never showed intranuclear aggregates in any of the HD cortices analyzed (**Figure 3H**). In contrast, Layer 3 IT, a resilient cell type for neuronal loss, had one of the largest proportion of cells with intranuclear aggregates (4%) (**Figure 3H**). Overall, when correlating intranuclear aggregation with neuronal loss, three patterns were observed: some cell types (including VIP, LAMP5, and L5 ET) appeared resistant to both intranuclear aggregation and neuronal loss; others (including SST and L5–6 NP) showed reduced numbers in HD despite being largely free of intranuclear aggregates; and a third group (including L3 IT, L5 IT, L6 IT, and L6b) had correlated levels of intranuclear aggregation and neuronal loss (**Figure 3I**). These findings indicate that intranuclear aggregation alone does not predict which cell types are more vulnerable to depletion in the HD cortex.

### Large somatic CAG expansions correlate with the loss of excitatory neuronal subtypes

While large somatic repeat expansion was identified in HD brain^32^ shortly after identification of the *HTT* gene^33^, it remains unclear which cell types undergo such large somatic repeat expansions, particularly in the human cortex. To estimate the number of CAG repeats in individual cells of human brain tissue, we developed a MERFISH-compatible single-molecule FISH (smFISH) approach reporting on repeat length (**Figure 4A**). The method combines, in an additional cycle of hybridization, 56 oligonucleotide probes targeting 5’ *HTT* exonic sequences detected in a near infra-red channel, with a CAG-probe targeting a 13-repeat CAG sequence that was amplified via branching tree amplification^34^ detected in a far-red channel (**Figure S9A-C**). For each *HTT* transcript, detected as a smFISH punctum, the brightness of the colocalizing CAG-probe was quantified and then normalized by the brightness of the *HTT* probes (**Figure 4B and S9D**).

**Figure 4:**
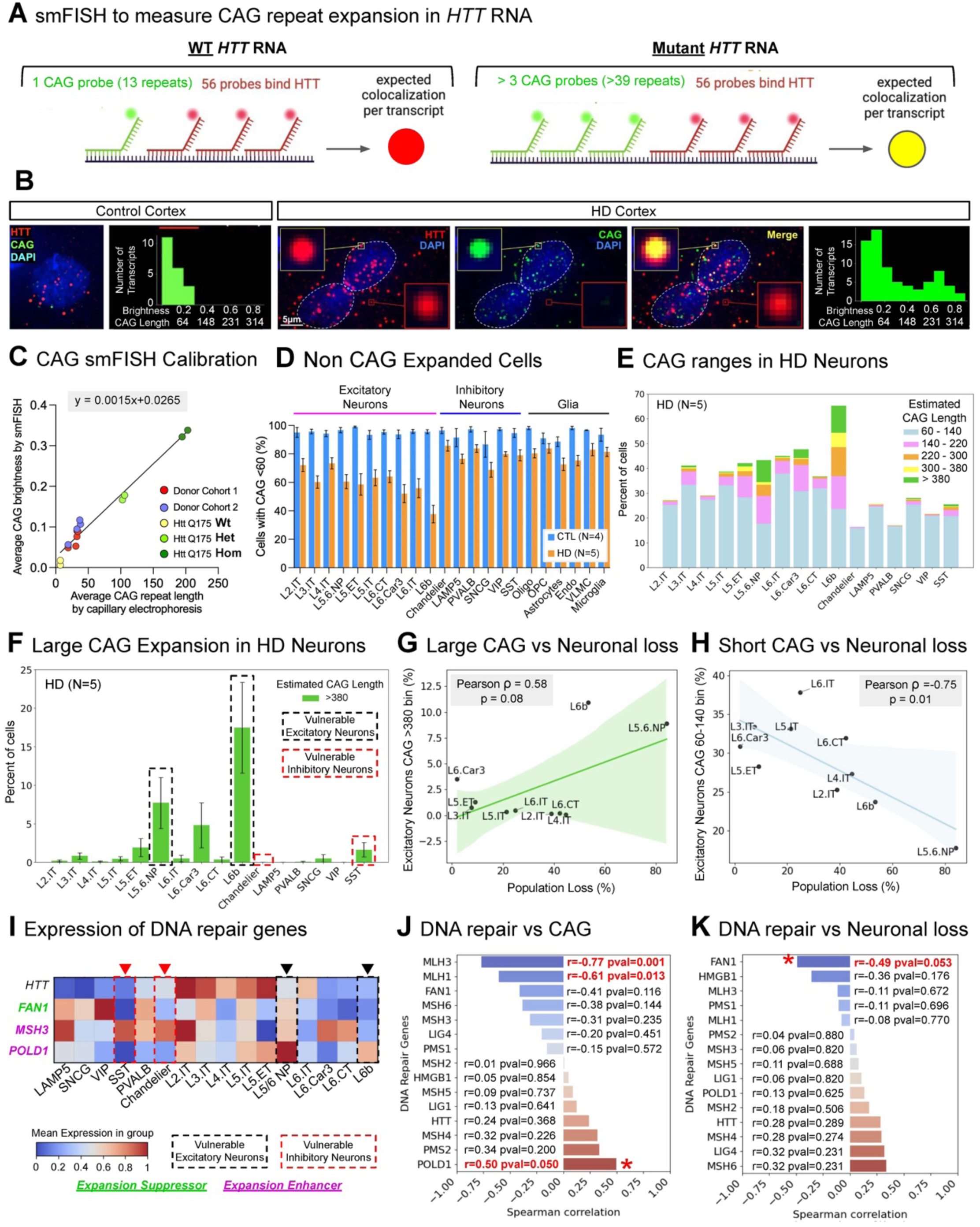
Excitatory neuron loss was associated with very large (>380) somatic CAG expansions. **(A)** Schematic cartoon of the smFISH probe design to quantify CAG repeats in HTT RNA. 56 probes are targeting the exons of HTT RNA and read in ‘red’. One unique CAG probe targeting 13 CAG repeat is read in ‘green’. Brightness of CAG probe colocalizing with HTT transcripts would be proportional to the CAG track length, we expect a constant amount of ‘red’ signal in HTT transcript while the CAG ‘green’ signal will increase as somatic repeat expansion occurs. (**B**) Representative images of *HTT* transcripts (red) and CAG probe (green) in Control cortex (Left, with unimodal distribution of CAG probe brightness) and HD cortex (Right, Bimodal distribution of CAG probe). In HD cortex (right), two HTT transcripts with high (top) and low (bottom) CAG signal are highlighted. DAPI in blue. (**C**) Correlation between mean CAG in *HTT* transcripts by smFISH and as measured by capillarity electrophoresis in human and mice cortices. (**D**) Percentage of cell with >60 estimated CAG repeats in *HTT* in control and HD cortices. (**E**) Percentage of cells that have 60-140, 140-220, 220-300,300-380 or more than 380 estimated CAG repeats in *HTT* in HD cortices. (**F**) Percentage of cells that have more than 380 estimated CAG repeats in *HTT* in HD cortices. (**G**) Correlation between the percentage of cells with more than 380 estimated CAG repeats in *HTT* in each neuron with the percentage of cells lost within that cluster in HD cortex. (**H**) Correlation between the percentage of cells estimated 60-140 CAG repeats in *HTT* in each neuron with the percentage of cells lost within that cluster in HD cortex. (**I**) Heatmap of expression of DNA repair genes involved in HD expressed in neuronal clusters from MERFISH. Magenta genes are known to be CAG expansion enhancer and green genes expansion suppressor^12^. (**J**) Spearman correlation for DNA repair genes expression levels measured by MERFISH in each neuronal cluster and percentage of cells with more than 140 estimated CAG repeats in *HTT*. (**K**) Spearman correlation for DNA repair genes expression levels measured by MERFISH in each neuronal cluster and percentage of cells lost within each cluster.

To convert the normalized CAG brightness into an estimated CAG repeat length, we supplemented the human cortical samples with measurements of 1) isogenic Embryonic Stem Cells (ESCs) harboring heterozygous *HTT* alleles with known 30 and 81 CAG repeats and 2) mouse models harboring 180–220 CAG repeat expansions in the murine *Htt* gene (HttQ175). The average CAG brightness per transcript distinguished between the expanded and non-expanded alleles (**Figure S10A-D**), and scaled linearly with the average number of CAG repeats per sample spanning from 7 to 200 CAG repeats (**Figure 4C** and **S10E-F**). We hence calculated a linear calibration function for converting between CAG brightness and estimated CAG repeat length. At a single-molecule level, heterozygous mice containing a non-expanded allele of 7 CAG repeats and an expanded allele with 205 CAG repeats displayed a bimodal distribution for the estimated CAG repeat length per *Htt* transcript with one peak centered at 7 CAG repeats and one centered at 205 CAG repeats (**Figure S10G**). In contrast, the transcripts from wild-type and homozygous HttQ175 mice had unimodal distributions centered at 7 and 200 CAG repeats, respectively (**Figure S10G**). Importantly, the mouse data allowed for quantifying the accuracy of the CAG repeat length as a function of number of molecules that were quantified and averaged: the standard deviation for estimated of CAG repeat length was 67 CAGs when averaging 10 molecules, but reduced to 21 CAGs when averaging 100 molecules (**Figure S10H**).

When examining the *HTT* transcripts of individual cells in the human cortical samples we observed that some neurons in the HD patients had bimodal distributions of estimated CAG lengths across *HTT* transcripts, similar to the heterozygous mouse models. The peaks of these distributions likely correspond to transcripts from the non-expanded and the expanded *HTT* alleles (**Figure 4B** – cell with an estimated CAG length of ∼250 in the mutant allele). In contrast, the *HTT* transcripts in the control patient samples have unimodal distributions (**Figure 4B**). Since, on average, each neuron contains 15 *HTT* transcripts, we estimate that the accuracy of our single neuron measurements of CAG expansion is ±55 CAG repeats.

In single cells, large somatic expansions (>140±55 estimated CAG repeats) were detected exclusively in HD cortical cells compared to controls (**Figure 4D** and **S11CS11C-G**), with the highest frequencies observed in L5–6 NP and L6b excitatory neurons (**Figure 4E** and **S16**). In these neuronal types (L5.6 NP and L6b), we also observed very large expansions (>380±55 estimated repeats) (**Figure 4F**). Overall, the proportion of cells with very large expansions (>380±55 estimated repeats) correlated with selective vulnerability among excitatory neuronal types (**Figure 4G**, ρ = 0.58, p = 0.08). In line with this, excitatory neuronal types lacking or with short somatic expansions (60-149±55 estimated repeats) correlated negatively with neuronal loss (ρ = -0.75, p = 0.01) (**Figure 4G**). For example, L5–6 NP and L6b neurons had the highest degree of somatic expansion and were the most depleted in HD cortex, while L5 ET and L3 IT did not present large somatic expansion and had comparable number of cells in HD cortex compared to controls. For inhibitory neurons, the correlation between somatic expansion and vulnerability was less pronounced (>380±55 estimated repeats, ρ = 0.33, p = 0.53 and 60-140±55 estimated repeats, ρ = -0.54, p = 0.26) (**Figure S11F-G**) suggesting that there might be additional factors contributing to their vulnerability.

We next investigated whether the expression of DNA mismatch repair genes is associated with the differential vulnerability of cortical neuronal subtypes to undergo somatic repeat expansion. We quantified the expression of HD related DNA repair machinery genes across neuronal populations and correlated these levels with the percentage of cells harboring large somatic repeat expansion (>140 estimated CAG repeats) (**Figure 4I-J** and **S12**). Expression of *POLD1*, a DNA polymerase whose action enhances somatic repeat expansion in mice^12^, was negatively correlated with somatic expansion (ρ = 0.50, p = 0.05) (**Figure 4J**). Conversely, expression of *FAN1* , a nuclease identified as a modifier of age at onset in HD^10^ and suppressor of somatic repeat expansion^12^, was negatively correlated with vulnerability (ρ = -0.49, p = 0.05) (**Figure 4K**). *FAN1* expression was highest in resilient inhibitory neurons including those expressing vasoactive intestinal peptide (VIP), while POLD1 was highest in the most vulnerable L5-6NP neurons (**Figure 4I**). This analysis was consistent with quantifications of gene expression based on single-nucleus RNA-seq^25^ (**Figure S12**). Interestingly, *MSH3*, a core component of the MutSβ mismatch-repair complex that is essential for somatic CAG repeat expansion in the HD striatum ^12,13^ did not correlate with long CAG repeat expansion in HD cortex (**Figure 4I-J**).

### Intranuclear aggregation is most prevalent at an intermediate level of somatic repeat expansion

To determine how somatic CAG repeat expansion relates to intranuclear HTT aggregation at single-cell level, we examined the co-occurrence of CAG expansions and nuclear inclusions. Cells with large somatic repeat expansions frequently—but not always— contained intranuclear *HTT* aggregates (**Figure 5A**). For example, Layer 5–6 Near-Projecting (L5–6 NP) neurons exhibited substantial somatic expansions despite the absence of detectable intranuclear inclusions, whereas nearly 40% of Layer 6b neurons with large expansions also harbored intranuclear *HTT* aggregates (**Figure 5B**). Across all cell types, cells containing intranuclear aggregates had significantly longer estimated CAG repeat lengths, with a mean of 149±55 repeats (**Figure 5C**). When stratifying the percent of cells with nuclear aggregates across different intervals of CAG repeat lengths, we noticed that aggregates appeared most prevalently at an intermediate level of somatic repeat expansion (220-300±55 repeats) before becoming less prevalent at most extreme lengths of repeat expansion (**Figure 5D**). This result was consistently observed across all HD cortex samples analyzed (**Figure S14A-B**).

**Figure 5:**
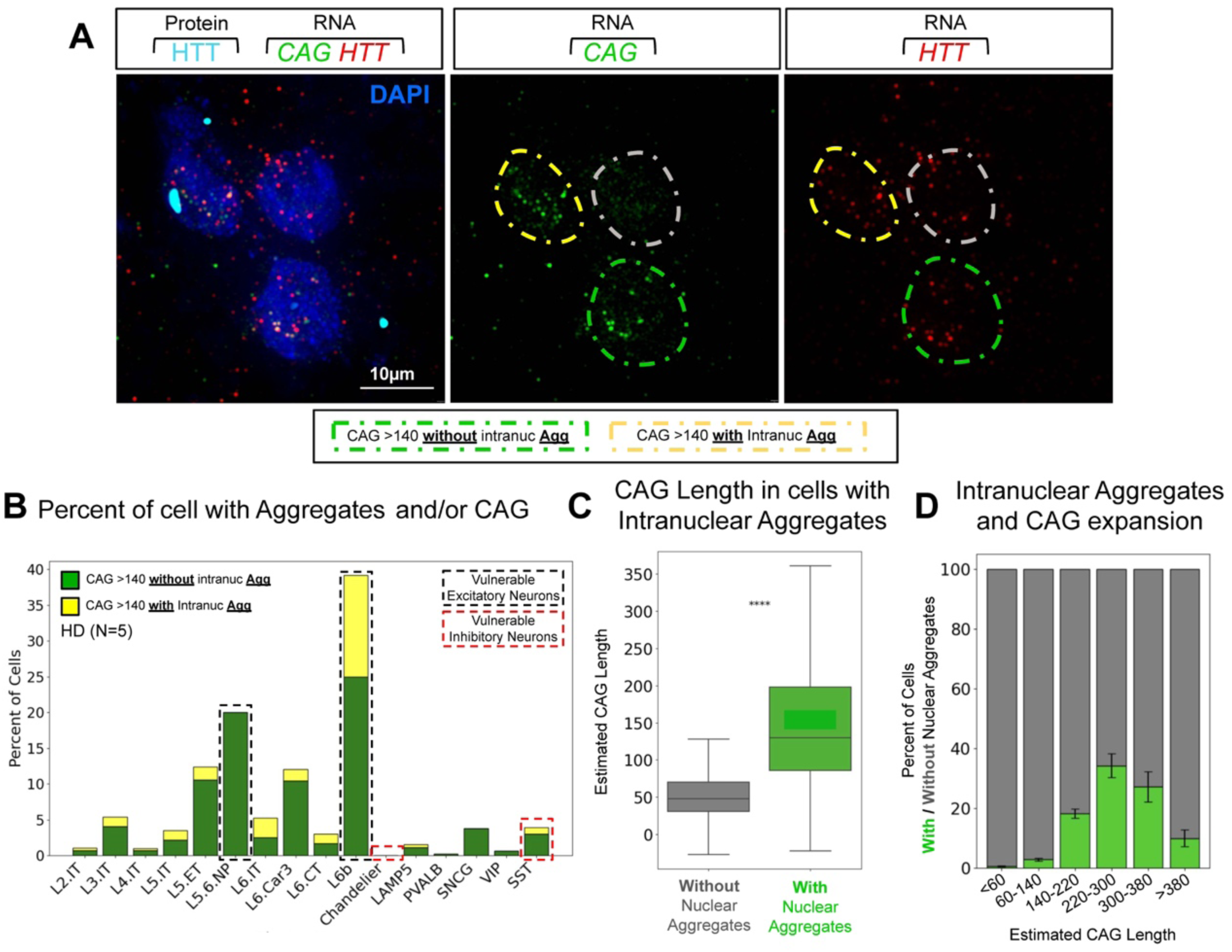
Intranuclear aggregation is most prevalent at an intermediate level of somatic repeat expansion. **(A)** Representative image of *HTT* transcripts (red), CAG probe (green) and HTT protein in cyan in HD cortex DAPI in blue. **(B)** Cells with aggregates with or without CAG expansion. (**C**) Estimated CAG length in HTT RNA in cells without or with intranuclear HTT aggregates. (**D**) Percentage of cells with and without intranuclear HTT aggregates across different estimated CAG length in HTT RNA per cell.

### Markers of DNA damage and chromatin silencing are increased in cells with somatic repeat expansion and HTT intranuclear aggregation

To further characterize the selective vulnerability of cell types with somatic repeat expansion and HTT aggregation we examined additional potential pathogenic marks including a DNA damage marker and histone modifications associated with open or closed chromatin. Because double stranded DNA damage has been reported to increase in HD^9^ and can lead to transcriptional arrest^35^ , we quantified γH2AX, a marker of double-stranded DNA breaks. γH2AX intensity was higher in cells containing intranuclear aggregates and rose with increasing CAG repeat length, peaking at an intermediate level of 220-300±55 repeats, similarly to the intranuclear aggregates (**Figure 6A-C**). The repressive epigenetic mark H3K9me3 was also elevated in cells with intranuclear aggregates and similarly was more frequent in neurons with an intermediate repeat length (220-300±55 repeats). In contrast, the mark of active chromatin H3K27ac remained largely unchanged (**Figure 6D-I**). Finally, we quantified lipofuscin, an auto fluorescent aggregate that accumulates progressively with cellular aging^36,37^. Lipofuscin granule numbers were significantly higher in neurons with HTT intranuclear aggregates and progressively increased with somatic repeat expansion (**Figure S13C-E**). Together, these findings suggest that DNA damage, chromatin silencing and aging-related metabolic burden accompany somatic repeat expansion and HTT aggregation and may contribute to their selective vulnerability.

**Figure 6:**
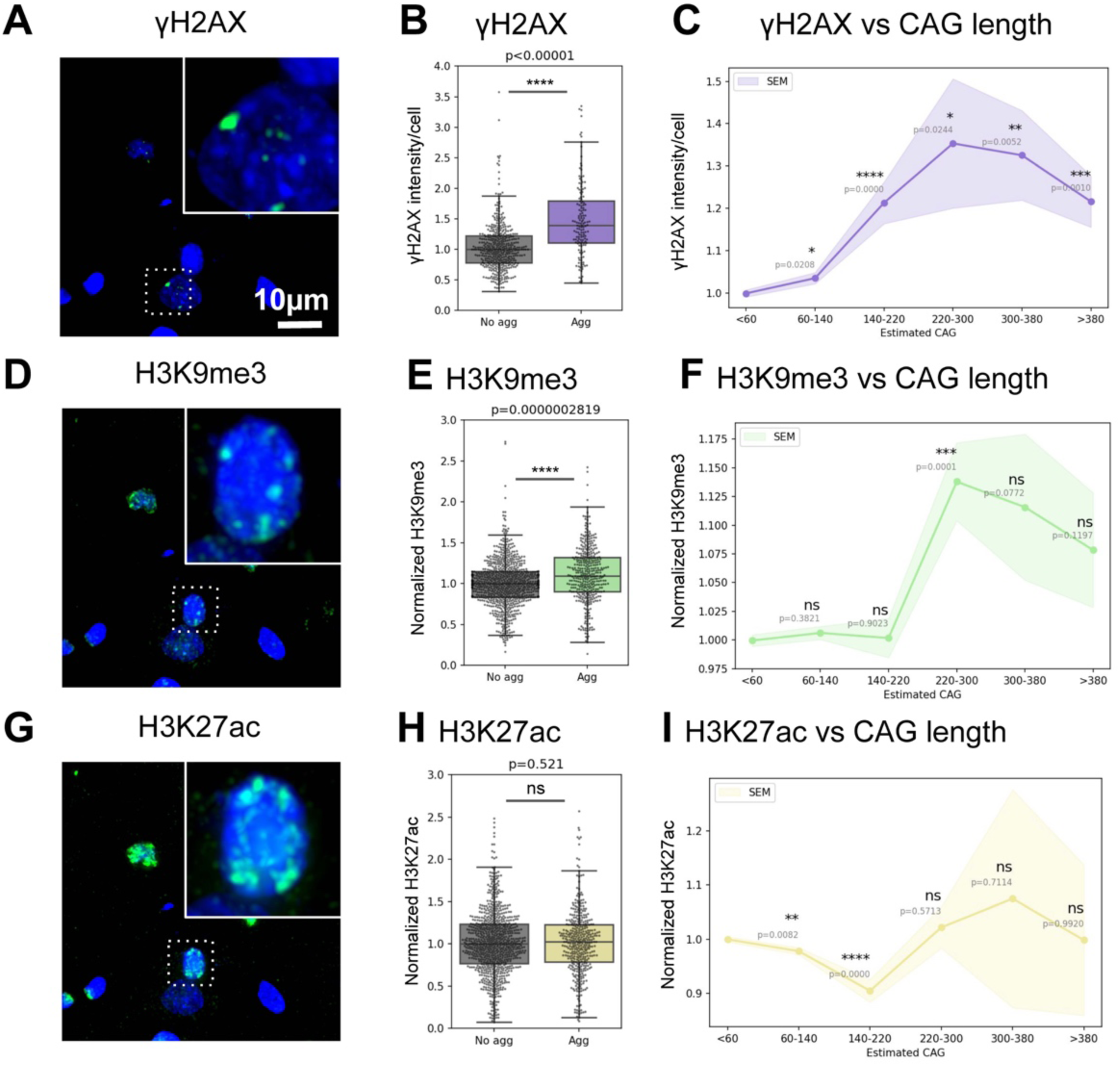
DNA damage and chromatin silencing are increased in cells with somatic repeat expansion and HTT intranuclear aggregation. **(A)** Representative image of γH2AX in HD cortex. **(B)** Quantification of γH2AX in excitatory neurons without or with intranuclear aggregation. **(C)** Quantification of γH2AX in excitatory neurons that have 60-140, 140-220, 220-300,300-380 or more than 380 estimated CAG repeats in *HTT* in HD cortices. **(D)** Representative image of H3K9me3 in HD cortex. **(E)** Quantification of H3K9me3 in excitatory neurons without or with intranuclear aggregation. **(F)** Quantification of H3K9me3 in excitatory neurons that have 60-140, 140-220, 220-300,300-380 or more than 380 estimated CAG repeats in *HTT* in HD cortices. **(G)** Representative image of H3K27ac in HD cortex. **(H)** Quantification of H3K27ac in excitatory neurons without or with intranuclear aggregation. **(I)** Quantification of H3K27ac in excitatory neurons that have 60-140, 140-220, 220-300,300-380 or more than 380 estimated CAG repeats in *HTT* in HD cortices.

### Transcriptional dysregulation is most strongly associated with nuclear huntingtin aggregation, rather than somatic CAG repeat expansion

To disentangle how somatic repeat expansion and HTT aggregation correlate to transcriptional changes, we performed a series of focused pairwise comparisons. First, transcriptional differences in neurons with or without large somatic expansions within the same samples (>280±55 vs <60±55 estimated repeats) included downregulation of *HTT* itself, and upregulation of genes enriched for developmental transcription factors (*TBX5, POU4F2, HAND2, DACH1, SIX1,* and *HOXD9*) (**Figure S13**). The magnitude of these changes increased with repeat length, reaching ∼1.5-2 fold by >350±55 CAG repeats (**Figure 7A-B and S13**). To better control for cell type effects and spatial context, we compared each neuron with a large expansion (>280±55 estimated CAGs) to its nearest non-expanded neighbor of the same neuronal type (<60±55 estimated CAGs) (**Figure S7C-D**). In this comparison, we detected overall modest transcriptional changes among the genes measured. The most upregulated genes included *CBX8* (a component of Polycomb Repressive Complex 1) and the transcription factor *HAND2*, each showing ∼1.5-2-fold increases in RNA levels (**Figure 7D**).

**Figure 7:**
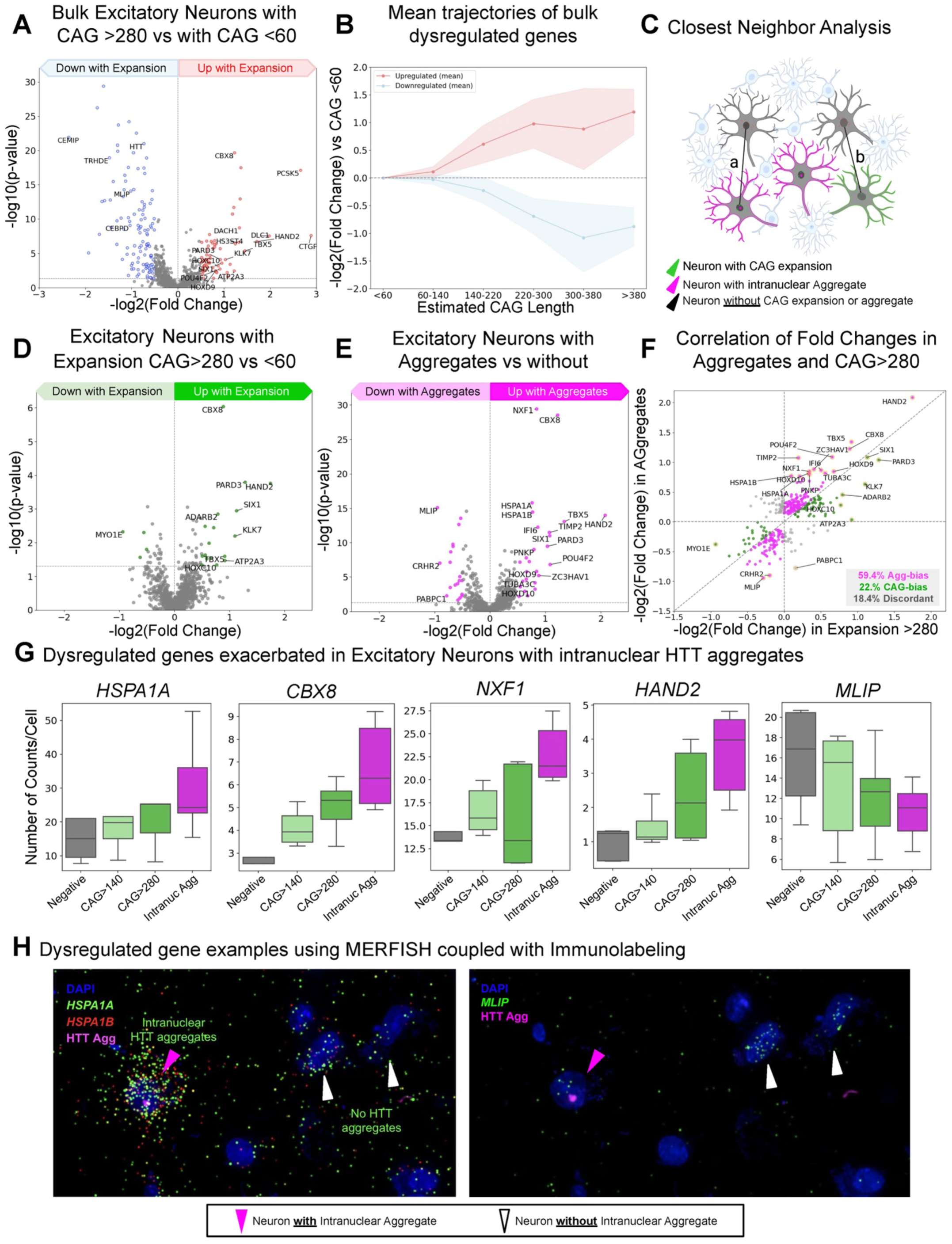
Transcriptional changes are more closely associated with huntingtin inclusions than with large repeat expansions. **(A)** Schematic cartoon of the closest neighbor analysis used to a series of pairwise comparisons between individual cells. For example, in (B), we compared neurons with large somatic expansions (>140 estimated CAG repeats) but no intranuclear HTT aggregates to their nearest neighbors of the same cell type that lacked both somatic expansion and aggregates. **(B)** Volcano plot of statistical significance versus the log2 fold change for gene expression in neurons with large somatic expansions (>140 estimated CAG repeats) but no intranuclear HTT aggregates compared to their nearest neighbors of the same cell type that lacked both somatic expansion and aggregates. **(C)** Volcano plot of statistical significance versus the log2 fold change for gene expression in neurons with large somatic expansions (>280 estimated CAG repeats) but no intranuclear HTT aggregates compared to their nearest neighbors of the same cell type that lacked both somatic expansion and aggregates. **(D)** Volcano plot of statistical significance versus the log2 fold change for gene expression in neurons with intranuclear aggregates compared to their nearest neighbors of the same cell type that lacked intranuclear aggregates. **(E)** Plot of log2 fold change for gene expression in neurons with intranuclear aggregates versus the log2 fold change for gene expression in neurons with large somatic expansions (>140 estimated CAG repeats). **(F)** Plot of log2 fold change for gene expression in neurons with intranuclear aggregates versus the log2 fold change for gene expression in neurons with large somatic expansions (>280 estimated CAG repeats). **(G)** Number of *HSPA1A, CBX8, NXF1, HAND2* and *MLIP* RNAs in neurons without expansion or intranuclear aggregates (Negative), neurons with large expansions (>140 and >280 estimated CAG repeats) and neurons with intranuclear aggregates. **(H)** Representative image of neurons with (magenta arrowhead) and without (white arrowhead) depicting *HSPA1A, HAPA1B* and *MLIP* MERFISH decoded transcripts.

In parallel, we compared cells that had intranuclear HTT aggregates to their closest aggregate-free neighbors of the same cell type. In this case, we observed more prominent transcriptional changes, with *CBX8* and *HAND2* among the most upregulated genes by approximately 5.3-fold (**Figure 7E**). To dissect the contribution of HTT intranuclear aggregation, we correlated the fold changes of differentially expressed genes in aggregate-bearing cells and in cells with large repeat expansions (280±55 CAGs). The majority of genes exhibited consistent upregulation or downregulation across both states (only 18.4% were discordant), despite only 19.6% of neurons with intranuclear HTT aggregation having >280±55 CAG expansion (**Figure 7E-F**). Remarkably, the presence of intranuclear aggregates tended to amplify transcriptional changes: for most differentially expressed genes, the magnitude of the fold change was greater in aggregate-bearing cells (59.4%), whereas fewer genes (22%) showed stronger changes in expanded cells (**Figure 7E-F**). The presence of intranuclear aggregates enhanced the upregulation of *HAND2*, as well as *HSPA1A* and *HSPA1B* (**Figure 7E-H**), members of the HSP70 family of molecular chaperones known to respond to protein aggregation induced stress^38^. We also detected increased expression of *NXF1*, a nuclear export factor involved in mRNA transport, and decreased *MLIP* (muscular Lamin A-interacting protein) (**Figure 7E-H**), consistent with prior evidence that aggregate-bearing cells experience RNA trafficking and nuclear envelope stress^9^.

## DISCUSSION

Multiple mechanisms have been proposed to contribute to HD pathogenesis including aberrations in autophagy and proteasome function, transcriptional dysregulation, disruption of nuclear import and export, increased DNA damage, impaired axonal transport, synaptic dysfunction, excitotoxicity, and metabolic deficits^1,39^. Among these, the two most intensively studied and mechanistically linked processes are huntingtin aggregation and somatic CAG repeat expansion^1,39^. Yet how these processes relate to selective neuronal loss in the human HD brain has remained unresolved because prior methods lacked the molecular and spatial resolution and the multimodal capabilities to disentangle them. Recent sequencing efforts from the Heintz^16,17^ and McCaroll^15^ groups initiated efforts to address this challenge. However, quantifying all the essential modalities, *in situ,* within intact sections of diseased human brains, including ***1)*** fine definition of cell types, ***2)*** the number and size of huntingtin inclusions, and ***3)*** the extent of somatic CAG expansion (particularly for very large expansions), cannot be currently explored by their sequencing approaches. By developing our Multimodal MERFISH approach, we overcame these limitations to determine that cortical pathology is not uniform but heterogeneous, localizing to specific populations of excitatory and inhibitory neurons, each with different vulnerabilities for HTT aggregation and somatic expansion.

### Large somatic repeat expansions are associated with the loss of specific excitatory neurons

A major technological advancement introduced here is the extension of the multimodal MERFISH to provide the first imaging-based approach for estimating somatic repeat expansion at single-cell resolution within intact human tissue. Our expansion estimates exhibit a per-cell variability of ±55 repeats, which likely reflects hybridization fluctuations arising from probes targeting a 13-CAG motif. While the precision is lower compared to sequencing approaches, this hybridization-based method is particularly suited for characterizing the larger spectrum of somatic expansion (i.e. >200 repeats), complementing sequencing-based assays which are biased towards the smaller spectrum of repeat length. Specifically, PCR-based or short-read sequencing methods systematically underrepresent long alleles due to size-dependent amplification bias and read-length limits (typically ∼100–150 bp)^16,17,40^. Recent long-read sequencing approaches only partly mitigate this bias^15,40^, still suffering from preferential loading of shorter molecules compared to the highly expanded alleles. In contrast, our assay does not rely on enzymatic amplification and can therefore detect very long expansions without distorting the distribution. This short-scale bias could obscure the full mechanism of the disease: for instance, recent short-read sequences of neuronal populations from the cortex and striatum of HD patients ^15,16^ suggested that somatic instability is widespread, yet not highly predictive of cell death. In contrast, recent work combining snRNA-seq with long-read sequencing of *HTT* detected highly expanded alleles (>400 CAGs) in the most vulnerable medium spiny neurons of the striatum. Our measurements integrate these perspectives: we observed a broad distribution of smaller somatic expansions (<140 ± 55 CAGs) across many cortical excitatory neuronal types, but subpopulations of neurons with increasingly higher somatic expansion have increasingly higher neuronal loss.

### Differential expression of the HD-related DNA repair genes across vulnerable neuronal populations

Cortical excitatory neuron populations share broad transcriptional similarity across their transcriptomes, yet the differential expression of specific genes might explain their vulnerability to cell death and their susceptibility to increased somatic repeat expansion. We specifically searched across the HD related DNA repair factors and found that *POLD1*, a DNA polymerase that promotes repeat instability *in vivo*^12^, was markedly enriched in the most affected excitatory populations—L5–6 NP and L6b neurons—which also exhibit the largest somatic expansions. In contrast, *FAN1*, a nuclease known to suppress CAG expansions *in vivo*^12^ and whose increased expression delays onset in humans, was highest in the most resilient excitatory subtypes. In the striatum, elevated *MSH2* and *MSH3* levels in MSNs have been proposed to explain their high expansion burden and early loss^17^. In contrast, in vulnerable cortical neurons we did not see this *MSH3* enrichment. Considering that *MSH3* is currently a prominent therapeutical target^14,41^, it becomes critical to determine whether its effect in cortical neurons mirrors its effects in striatal neurons.

### Connecting somatic CAG repeat expansion, aggregation, and transcriptional dysregulation

Previous studies using bulk or dissociated-cell sequencing reported transcriptional signatures linked to repeat instability but could not separate the influence of expansions from that huntingtin aggregation. We found that neurons harboring large expansions showed overall modest transcriptional changes, including upregulation of developmental transcription factors such as *HAND2* and *HOXD9*. Similar transcription factor activation and loss-of-identity signatures have been reported in HD by bulk^42^ and single-cell sequencing^15^ studies. Neurons that carry intranuclear aggregates exhibit broader transcriptional changes, including upregulation of *HAND2* and other developmental transcription factors which are further amplified, and induction of *HSPA1A* and *HSPA1B*, members of the HSP70 family of molecular chaperones that are well known to respond to protein aggregation stress^38^.

### Implications for other neurodegenerative disorders

By preserving spatial architecture and enabling simultaneous measurement of genomic instability, protein aggregation, and transcriptional state at single-cell resolution, our multimodal MERFISH approach provides a powerful framework for resolving how these processes interact within intact human tissue. We expect this approach to be extendable to other pathologies with repeat expansions —including *ATXN*s CAG repeat expansions in spinocerebellar ataxias, *FMR1* CGG repeat expansions in fragile X–associated disorders, *FXN* GAA repeat expansions in Friedreich’s ataxia, and the *C9ORF72* G_4_C_2_ hexanucleotide repeat expansion in ALS/FTD^43^. By visualizing repeat instability, aggregate formation, and transcriptional remodeling within the same cells, multimodal MERFISH offers a scalable platform to untangle mixed pathologies and reveal how genomic and proteomic stressors shape selective vulnerability and resilience in the human brain.

## RESOURCE AVAILABILITY

### Lead contact

Further information and requests for resources and reagents should be directed to lead contacts Bogdan Bintu and Don W. Cleveland.

### Materials availability

Oligonucleotide probe sequences used for Multimodal MERFISH imaging can be found in Tables S1. Templates and reagents for making these probes can be purchased from commercial sources.

### Data and code availability

All Multimodal data reported in this paper are in the process of being submitted to Dryad.

Analysis code used for MERFISH probe library design, MERFISH image decoding, and data quantification is available at: https://github.com/BogdanBintu/NMERFISH and https://github.com/deprekate/MERMAKE/

Analysis code for downstream data processing is available at: https://github.com/oleolatz/HDMMERFISH

Any additional information required to reanalyze the data reported in this paper is available from the lead contacts upon request.

## MATERIALS and METHODS

### Autopsy materials

Human brain samples were obtained in collaboration from The Netherlands Brain Bank (NBB), Netherlands Institute for Neuroscience, Amsterdam (open access: www.brainbank.nl), and from the University of Washington UW Biorepository (UWA) and Integrated Neuropathology (BRaIN) Laboratory, which is supported by the Alzheimer’s Disease Research Center (AG066509) and the Adult Changes in Thought Study (AG006781). Information on patients can be found in **Table S2**. All materials have been removed of any patient identifiers. All Material has been collected from donors for or from whom a written informed consent for a brain autopsy and the use of the material and clinical information for research purposes had been obtained by the NBB and UWA.

### Mice

*Htt*Q175 mice were described earlier^44,45^ and were provided by CHDI foundation (CHDI-81003003) and maintained on a C57BL/6 background. Males and females heterozygous for the expanded allele were bred together to obtain a colony of wild- type, heterozygous and homozygous animals; CAG repeats were maintained between 160-220. Maintenance and experimental procedures were approved by the Institutional Animal Care (IACUC number: S00225) and Use Committee of the University of California, San Diego.

### Cell lines

Human IsoHD lines 30Q, 45Q, and 81Q were a kind gift from Mahmoud A. Pouladi and were fully characterized previouly^46^. Cells were plated onto Matrigel (Corning, cat. no. 354277) coated plates in Essential 8 Medium (E8; Thermofisher Scientific, cat. no. A1517001) and 10 μM ROCK inhibitor (RI; Y-27632; Selleckchem, cat. no. S1049). Cells were maintained in an incubator at 37 °C with 5% CO2 and fed every 1–2 days as needed. Cells were split using EDTA (0.5 mM; Life Technologies, cat. no. 15575020) for routine passaging. Medium was supplemented with 10 μM RI to promote survival during passaging.

### Multimodal Spatial Transcriptomics in human postmortem cortical samples

To achieve such multimodal, high-resolution, single-molecule imaging in human post-mortem brain samples, we introduced five key modifications to conventional MERFISH protocols (**Figure S1**): (1) anchoring of encoding probes to the tissue to improve signal stability across multiple rounds of imaging, (2) incorporation of gene-specific adaptors to enable seamless integration between smFISH and MERFISH modalities, (3) reduction of autofluorescence specific to human postmortem samples, (4) integration with protein immunolabeling to allow joint detection of RNA and protein, and (5) inclusion of a targeted strategy to measure CAG repeat length in *HTT* RNA at the single-molecule level.

First, each encoding probe was synthesized with an acrydite modification to enable covalent incorporation into a thin polyacrylamide gel that was cast directly onto the tissue sample, ensuring long-term probe retention and probe stability even if the target mRNA molecules were degraded. Indeed, this anchoring strategy allowed for extended rounds of hybridization and imaging while maintaining sample integrity. Correspondingly, to ensure such stability each of our imaging experiments reported here began and ended with imaging of *NRGN* and *GAD1* RNAs via smFISH, from which we determined that there was 94.7% signal retention after 72 rounds (**Figure S1**). Second,1128 gene-specific adaptor probes were designed to enabled imaging of a 1128 gene library (∼50 encoding probes per gene x 1128 genes = 56,400 unique encoding probes total) by single molecule fluorescence in situ hybridization (smFISH) imaging. This design allowed a flexible transition between smFISH and MERFISH approaches by incorporating the combinatorial code at the level of adaptor and readout hybridization rounds, rather than embedding the MERFISH barcodes directly into the encoding probes (**Figure S1**). Third, a major challenge in applying MERFISH to aged human brain tissue was the high level of tissue autofluorescence. To address this, we implemented extensive photobleaching prior to imaging. However, significant autofluorescence persisted in the 488 nm and 546 nm channels. To overcome this limitation, we strategically restricted all RNA and protein imaging to the 647 nm and 750 nm channels, where autofluorescence was minimal—an essential modification that enabled high-contrast single-molecule detection in human postmortem samples (**Figure S1**). Fourth, importantly, in contrast with commercial MERFISH, we did not digest the tissue samples, preserving the ability to conduct immunofluorescence staining (**Figure 1C** and **S1**). Fifth and finally, as detailed below, we developed a smFISH approach to measure CAG repeat expansion in *HTT* RNA consisting in 56 probes targeting the exons of *HTT* RNA read in 750 nm and one unique CAG probe targeting 13 CAG repeat read in 647 nm staining (**Figure 1C**).

### MERFISH gene selection and probe library design and construction

We designed a panel of 1128 genes to be imaged with combinatorial barcoded imaging^66^ including 357 genes for cell typing selected based on the Allen’s Institute’s single nuclear RNA sequencing (snRNAseq) dataset for human brain, and other genes relevant for neurodegenerative diseases (see **Table S1**). The encoding probe set that we used contained ∼50 encoding probes per RNA=. Encoding probes were designed using a publicly available pipeline (https://github.com/bil022/ProbeDesigner). The principles mainly include the detection of off-targets based on genome-wide 17-mer off-target counts, 30-42 mers RNA/DNA probe sequences, GC content or melting temperature (Tm), and the avoidance of repeated regions. Probe Designer is implemented with three steps: 1) Build a 17-mer index based on reference genome (DNA) or genome annotation files (RNA). 2) Scan 17-mer counts for selected loci or genome sequences. 3) Filter and rank probe candidate based on pre-defined selection criteria. These probes were designed to have to enable serial smFISH and MERFISH. After validation, the MERFISH strategy using this encoding library and adaptors (see **Table S1**) was employed in subsequent experiments by pooling adaptors together in a combinatorial manner in each hybridization round.

### Overview of experimental system

The physical setup used for performing these experiments consists of several components: A custom-built fluorescence microscope was used to acquire images, while a custom-built fluidics system was used to automatically perform buffer exchanges on the microscope stage. Custom software was used to synchronize and control the various components, and to automate many experimental steps. Below is a description of each of these elements.

#### Microscope setup for Image acquisition

Image acquisition was performed using a custom-built microscope system^63,74^. The system was built around a Nikon Ti-U microscope body with a Nikon CFI Plan Apo Lambda 60x oil immersion Fluidics system configuration.

The fluidics system consisted of several main components: a pump, a set of valves connected in series, a flow chamber in which the sample was mounted, and tubing and connectors. This system allowed for 20–34 rounds of hybridization (depending on the number of valves and the number of spots reserved for special buffers). In experiments where the number of hybridization rounds exceeded the capacity of the flow system, we replaced the buffers with new ones via the following procedure:

1. The output from the valve system was directly connected to the waste collection vessel, bypassing the sample-containing chamber.
2. All valves were washed using 50% formamide in water and then 2X SCC.
3. The new set of buffers was introduced, and the chamber was reconnected to the flow system.
4. The experiment resumed for the next round of hybridization.

#### Software for controlling experimental components

All system components were controlled using custom-built software, based on: https://github.com/ZhuangLab/storm-control. This software package is composed of several main modules, which work in concert:

1. ‘‘Hal’’ is the software package used to control and synchronize all illumination and microscope components. We note that in some cases it will be necessary to write drivers for components, which are not included in this package. Hal is also used to define imaging parameters, such as illumination strength, sequence of stage and illumination operations during imaging (e.g., during a z-scan), exposure time etc. 2. ‘‘Steve’’ is a module used to take mosaic images (i.e., a composite image made up of many individual fields of view) and select regions for imaging in experiments.
2. ‘‘Kilroy’’ is the software used to control the fluidics components, and to define pre-programmed sequences of operations to be performed as sets (e.g., the set of operations undertaken when a new round of hybridization is performed).
3. ‘‘Dave’’ can issue commands to both Hal and Kilroy and is used to automate the performance of data collection by defining in advance a complete set of fluidics system and microscope operations, as well as the order and time-lag in which they are to be performed.

### Probe synthesis

Primary/encoding probes were amplified from the template library described in Supplementary Tables 1. The generation of the encoding probe sets were prepared from oligonucleotide pools, as described previously^63^. Briefly, we first used limited-cycle PCR to amplify the oligo pools (Twist Biosciences) for approximately 20 cycles. The reverse primer used in this step also introduced and acrydite anchor at the 5’ end of the probes. Then, the resulting product was purified via column purification and underwent further amplification and conversion to RNA by a high-yield in-vitro transcription reaction using T7 polymerase (NEB, E2040S). Subsequently, RNA products were converted to single strand DNA with Maxima Reverse Transcriptase enzyme (Thermo Scientific, EP0751), and then purified via alkaline hydrolysis (to remove RNA templates) and DNA oligo purification kits (Zymo Research D4060). All primers were purchased from Integrated DNA Technologies (IDT).

### Readout and adaptor probe preparation

All readout and adaptor probes were ordered from IDT (see **Table S1**) and were diluted directly from this stock.

### MERFISH sample preparation

Multimodal MERFISH samples were prepared as similarly to as described in^66^ with modifications: Briefly, 16-μm-thick tissue sections were pre-cleared by immersing into 70% (vol/vol) ethanol and then 5% SDS in PBS for 4 minutes. Then the tissues were preincubated with hybridization wash buffer (40% (vol/vol) formamide in 2X SSC) for ten minutes at room temperature. After preincubation, the coverslip was moved to a fresh 60 mm petri dish and added with 50 ul of encoding probe hybridization buffer (2X SSC, 50% (vol/vol) formamide, 10% (wt/vol) dextran sulfate, and a total concentration of 1 μg/μl encoding probes. The sample was placed in a humidified 47°C oven for 18 to 24 hours then washed with 40% (vol/vol) formamide in 2X SSC, 0.1% Tween 20 for 30 minutes at room temperature. To anchor the RNAs in place, the encoding-probe–hybridized samples were embedded in 4% polyacrylamide gel for 6h, post-fixed with 4% (vol/vol) paraformaldehyde in 2X SSC and washed with 2X SSC. Samples were photo-bleached for 6h using a MERSCOPE Photobleacher (cat #10100003, Vizgen). See **Table S1** for details of encoding library, adaptor and readout used.

### Sequential hybridization for sequential or combinatorial FISH imaging

Each round of hybridization consisted of the following general steps: 1) Flow in the hybridization buffer with a set of oligonucleotide probes specific to each round, as described below, 2) Incubate for 30-90 minutes at room temperature,3) Flow wash buffer, 4) Incubate for ∼200s, 5) Flow in the hybridization buffer with readout probes, 6) Incubate for 30 minutes at room temperature, 7) Flow wash buffer, 8) Incubate for ∼200 s and 9) Flow imaging buffer.

Imaging buffer was prepared as described previously^63^, and consisted of 60 mM Tris pH 8.0, 10% w/v glucose, 1% Glucose Oxidase Oxygen Scavenger Solution (containing ∼100 mg/mL Glucose Oxidase (Sigma-Aldrich, G2133) and a 1:3 dilution of catalase (Sigma-Aldrich, C3155)), 0.5 mg/mL 6-hydroxy-2,5,7,8-tetramethylchroman-2-carboxylic acid (Trolox; Sigma-Aldrich, 238813) and 50 mM Trolox Quinone (generated by UV irradiation of a Trolox solution) (Cordes et al., 2009; Rasnik et al., 2006). Trolox was dissolved in methanol before being added to the solution. After preparation, the imaging buffer was covered by a ∼0.5 cm thick layer of mineral oil to prevent exposure to oxygen.

The hybridization buffer and wash buffer were made up of 35% and 30% formamide in 2X SSC, respectively, with the hybridization buffer also containing 0.01% v/v Triton-X. The hybridization buffer was kept separately for each hybridization round and contained two (for human) or three (for mouse) sets of readout probes.

Before the next round of readout probe or adaptor probe hybridization, fluorescent signals from the readout probes in the current round were removed by flowing 100% formamide and then 2X SCC.

#### Immunofluorescence staining for Multimodal MERFISH

Each round of immunofluorescence staining consisted of the following general steps: 1) Flow in the blocking buffer (1.5% BSA with 0.3% Triton X-100 in 1X PBS buffer), 2) Incubate for 30 minutes at room temperature, 3) Flow in the primary antibody solution (0.3% Triton X-100 in 1X PBS) containing the primary antibodies (see Supplementary Table 1), 4) Incubate for 2 hours at room temperature, 5) Flow wash buffer(2X SCC) and incubate for ∼200s (3x), 6) Flow in the secondary antibody solution (0.3% Triton X-100 in 1X PBS) containing the secondary antibodies (see Supplementary Table 1), 7) Incubate for 1 hour at room temperature, 8) Flow wash buffer (2X SCC) and incubate for ∼200s (3x), and 9) Flow imaging buffer.

### Image acquisition

For each experiment, we constantly selected imaging SVZ and hippocampus brain regions in mice and the SVZ in human. After each round of hybridization, we acquired Z-stack images of each FOV in 3 or 4 colors: 647 nm and 750 nm illumination (or 560 nm, 647 nm, and 750) were used to acquire FISH images, and 405 nm illumination was used to image DAPI. Images were acquired in all channels before the stage was moved and images were acquired at a rate of ∼10 Hz.

### Overview of Multimodal MERFISH analysis pipeline

The image analysis pipeline used in this study was implemented in Python, and all the codes are available at: https://github.com/BogdanBintu/NMERFISH. The overall pipeline consists of the following steps:1. Identify and segment all imaged nuclei, 2. Fit 3D-Gaussians to all detected fluorescent spots in imaging channels used RNA imaging, as well as for DAPI, 3. Correct sample drift using DAPI.

#### Flat-field correction

Images were first corrected to remove variations in illumination present on the edges and corners of images (flat-field correction) by calculating the per-pixel median brightness across the first round of imaging, separately for each color. Each image was then inversely scaled by this median brightness.

#### Deconvolution

Next, the images were deconvoluted using the Wiener algorithm with a custom point spread function (PSF) calculated for the specific microscope used for imaging to reduce noise and enhance the discreteness of fluorescent spots. Finally, a high-pass filter was used with a blur sigma of 30 to further improve signal-to-noise ratio.

#### Localization of Fluorescent Spots

After pre-processing images, local maxima within a 1-pixel radius were identified and further filtered to enforce a brightness of at least 3600 and a correlation with the point spread function of 0.25. The remaining local maxima were used for decoding.

#### smFISH quantification

For quantification of transcripts imaged individually as smFISH targets, the fluorescent spots identified in the previous step were further filtered based on a minimum brightness relative to their local background. This threshold was selected separately for each color channel and manually tuned to remove noisy and dim fluorescence while keeping bright puncta.

#### MERFISH decoding

To resolve RNA molecules from the identified fluorescent spots across imaging rounds, spots were first clustered into groups across images within a range of 2 pixels after correcting for drift. Due to the MERFISH codebook designed with a hamming weight of 4, clusters with at least 3 spots were kept allowing for single-bit-error correction, while clusters with 1 or 2 spots were discarded. For each cluster, a brightness vector was constructed using the spot brightness in each round, then L2 normalized and compared with the codebook.

To filter noise from the resulting decoded transcripts, a score was calculated for each transcript based on three metrics: 1) the average brightness across the fluorescent spots, 2) the distance of the brightness vector to the nearest spot in the MERFISH codebook, and 3) the average distance of each fluorescent spot to the median of all spots. These metrics were combined into a single score by calculating the combined Fisher p-value against the distribution of each metric across all decoded transcripts. The distribution of scores for decoded transcripts with a blank barcode identity were then compared to the distribution of scores for decoded transcripts with a gene identity, and a score threshold was chosen which separates the peaks of each distribution.

#### Cell Segmentation

Cell boundaries were determined in 3 dimensions (3D) using Cellpose v294. First, for each z-slice image, a 2D segmentation mask was generated using the DAPI channel with Cellpose’s “dapi” model, after flat field correcting and deconvoluting the DAPI image as described in “Image pre-processing”. Cells were then linked between adjacent z-slices to create a 3D segmentation.

#### Assigning transcripts to cells

Transcripts identified by both smFISH and MERFISH were assigned to cells by first adjusting the coordinates of each molecule by the calculated drift between the images used for segmentation and those in which the RNA were imaged. Next, the transcript coordinates were rounded to the nearest integer and assigned to the cell based on the value of the segmentation mask at each transcript’s coordinates.

#### Protein Quantification

Flat field corrected and deconvolved antibody images were thresholded and then clustered to determine local aggregates and quantify their size and mean brightness. These aggregates were then associated to the corresponding cell using a similar procedure as for the RNA MERFISH data.

#### Single-molecule FISH (smFISH) to measure somatic CAG repeat expansion in *HTT* RNA

The method combines a set of 56 oligonucleotide probes targeting *HTT* exonic sequences, detected in a near-infrared channel (750 nm), with a CAG probe targeting a 13-repeat CAG sequence. The CAG probe signal was amplified via branching tree amplification^34^ detected in a far-red channel (647 nm).

Individual *HTT* RNA spots were identified based on fluorescence intensity and filtered using on a minimum brightness threshold relative to the local background. This threshold was manually tuned to remove noisy and dim fluorescence signals while preserving bright, well-defined puncta. For each detected *HTT* transcript, a Gaussian distriution was fitted to quantify the fluorescence brightness of the colocalizing CAG probe as detected by the camera. The CAG signal intensity was then normalized to the fluorescence intensity of the corresponding *HTT* punctum (**Figure S9D**).

To convert the normalized CAG brightness into an estimated CAG repeat length, a linear calibration function was generated. For each patient, were averaged all CAG imaging based measurements (the normalized CAG brightnesses across all detected *HTT* transcripts), and regressed against the average CAG repeat length measured in each allege from cDNA isolated from an adjacent tissue section by capillary electrophoresis (outsourced to Laragen/TransnetYX). A separate linear regression was calculated for each experimental batch to account for inter-experiment variability (**Figure S10D**), while maintaining the same internal control (HD4) across experiments.

To ensure reliable quantification, we further restricted analysis to cells containing at least five *HTT* transcripts, in which *<u>all HTT molecules detected were averaged.</u>* To estimate the mutant allele CAG length at single-cell level, stability of the wild-type allele was assumed. The mean estimated CAG repeat length per cell was therefore calculated as:

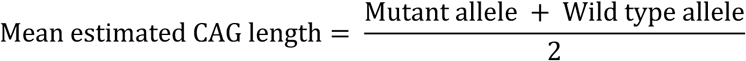

The accuracy of the CAG repeat length estimation as a function of number of molecules averaged was assessed using neonatal HttQ175 mice data. The standard deviation of the estimated CAG repeat length was 21 CAGs when averaging 100 molecules and 67 CAGs when averaging 10 molecules. Given that cortical human neurons contained, on average, 15 *HTT* transcripts per cell, we estimate the accuracy of single-neuron CAG length measurements to be approximately ±55 CAG repeats.

### Cell type quantification

Equal width columns with height defined by Layer 1/peripheral astrocytes to the white matter astrocytes are used to quantify cell numbers between control and HD cortices^28^.

### Statistics and graphs

Statistical analyses and graphs were performed using GraphPad Prism v8.0. For two-group analysis, Student’s t.test, ANOVA or the Mann-Whitney test was used, as determined by a normality test. All experiments include at least 3 biologically independent repeats; significance was set at *p<0.05, **p<0.01, ***p<0.001. Statistical significance was set at p< 0.05, and data are presented as mean ± SEM.

### Declaration of generative AI and AI-assisted technologies in the manuscript preparation process

During the preparation of this work the authors used ChatGPT in order to rephrase text or improve grammar. After using this tool, the authors reviewed and edited the content as needed and take full responsibility for the content of the published article.

## Acknowledgments

We would like to thank the families and patients of those affected by HD for their critical contributions to this research. This work was supported by CHDI (<u>A-20106).</u> Additional support was provided from the Hereditary Disease Foundation (HDF) to B.B., Huntington Disease Society of America (HDSA) to S.V-S and DP5-OD030878 to B.B.). OA-G was supported by the Center for Neurological Studies. R.M. was supported by postdoctoral fellowships from the HDF and the Bev Harting Huntington’s Disease Foundation and HDSA and K99/R00 (1K99AG090813-01). We would like to thank Stephen Moore for calculating the signal retention after 72 MERFISH cycles. We would like to thank the Netherlands Brain Bank and the University of Washington UW Biorepository and Integrated Neuropathology (BRaIN) Laboratory for supplying the human brain tissue for our study.

## Supplementary Figures

**Supplementary Figure S1:**
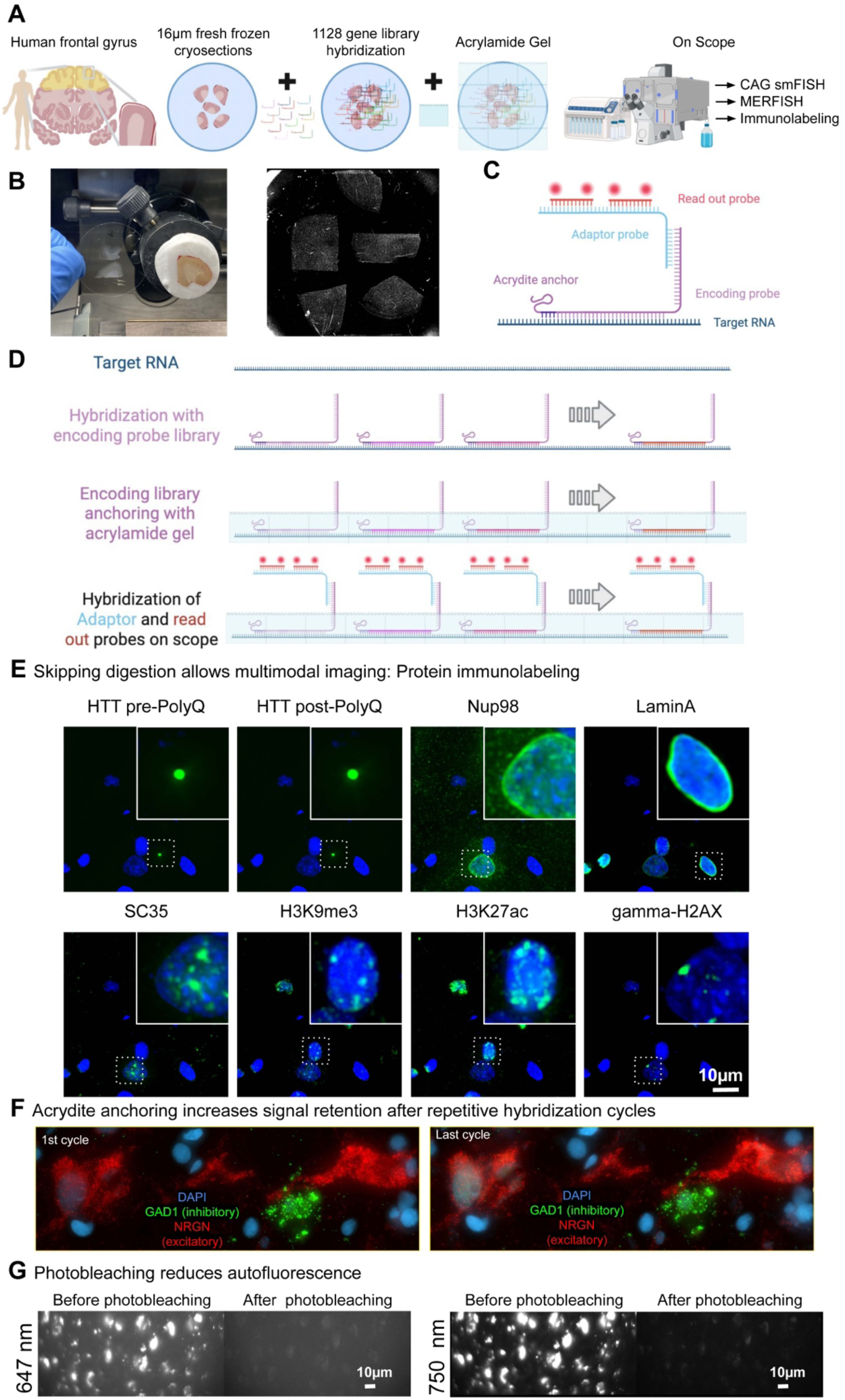
Multimodal MERFISH overview. (**A**) Overview of sample preparation, imaging and data processing workflow. Human cortical samples are cryosectioned at 16µm thickness, hybridized with a library encoding for 1128 RNAs, embedded in a polyacrylamide gel, and photobleached. Sequential rounds of smFISH, MERFISH and immunolabelling are performed on scope. (**B**) Cryosectioning of fresh frozen human brain cortex. (**C** and **D**) Illustration of the primary DNA probes used for single molecule FISH (smFISH)/MERFISH/Multimodal MERFISH experiments. Each probe is designed to target a complementary mRNA sequence (40 bases) and contains one unique barcode (20 bases) per gene detected by hybridization with adaptor and readout probes. The probes were synthesized with an acrydite anchor modification on the 5’ end to allow for a covalent incorporation into a thin acrylamide gel cast on top of the sample after hybridization. (**E**) Representative images of sequential immunofluorescence and imaging for HTT, NUP98, Lamin A, SC35, H3K9me3, H3K27ac and γH2AX. (**F**) Representative images of smFISH before and after 1128 MERFISH experiment from the human frontal cortex. (**G**) Representative images of autofluorescence recorded at 647 nm before and after photobleaching and 750 nm before and after photobleaching from the human frontal cortex. (**F**) Representative images of autofluorescence recorded at 647 nm before and after photobleaching and 750 nm before and after photobleaching from the human frontal cortex.

**Supplementary Figure S2:**
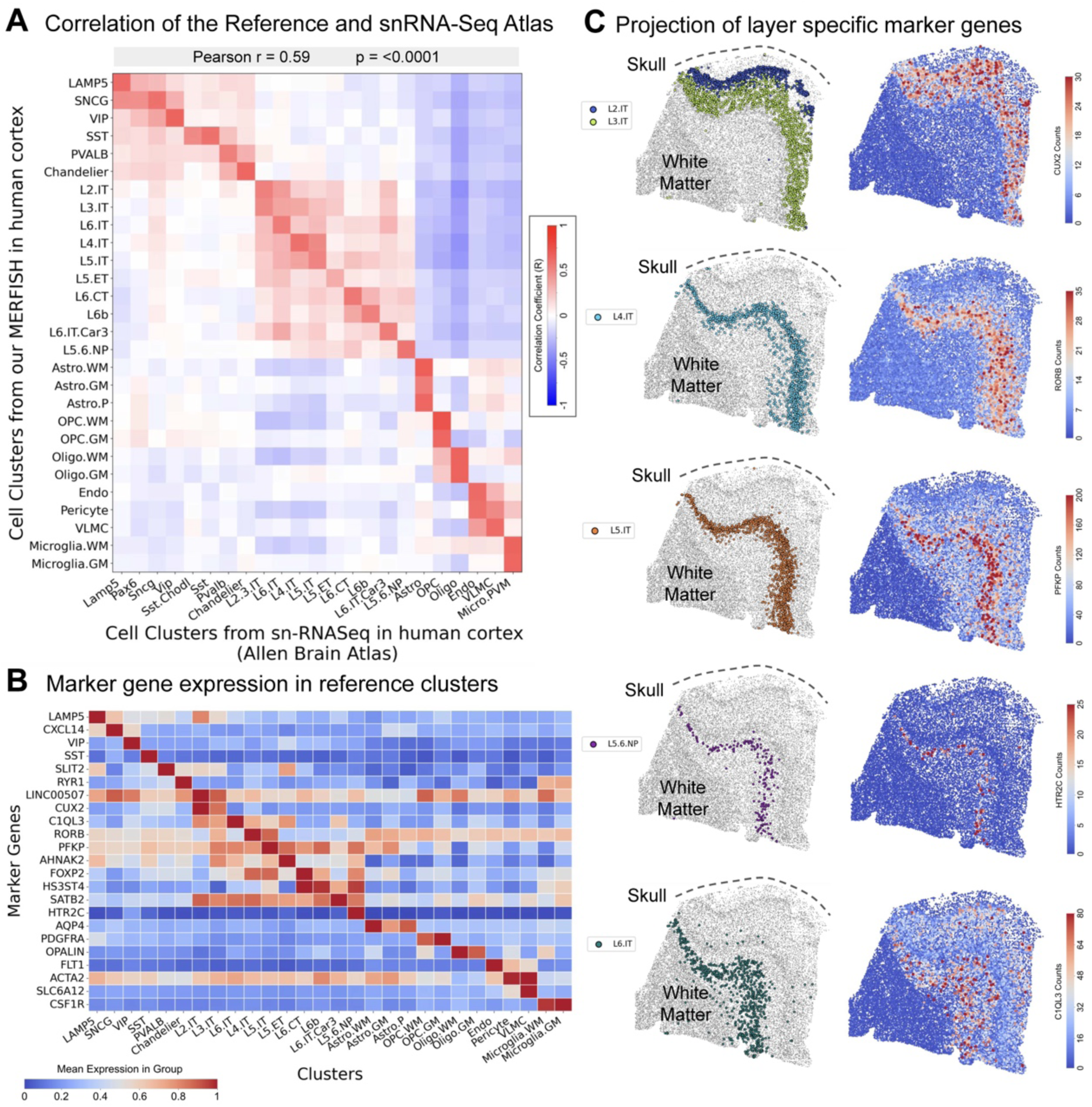
MERFISH clustering of 188 gene measurements aligns with snRNA-seq. (**A**) A Pearson correlation heatmap comparing the major cell type clusters in MM-188 genes MERFISH (Control 4 with 29,456 cells) and the cell clusters from single nuclei RNA sequencing in human cortex (Allen Brain Atlas). (**B**) Heatmap shows normalized mean expression level for cell-type-specific marker genes across in the human control cortex. (**C**) Spatial projection of cell-type-specific marker genes in Reference Control 4 cortex.

**Supplementary Figure S3:**
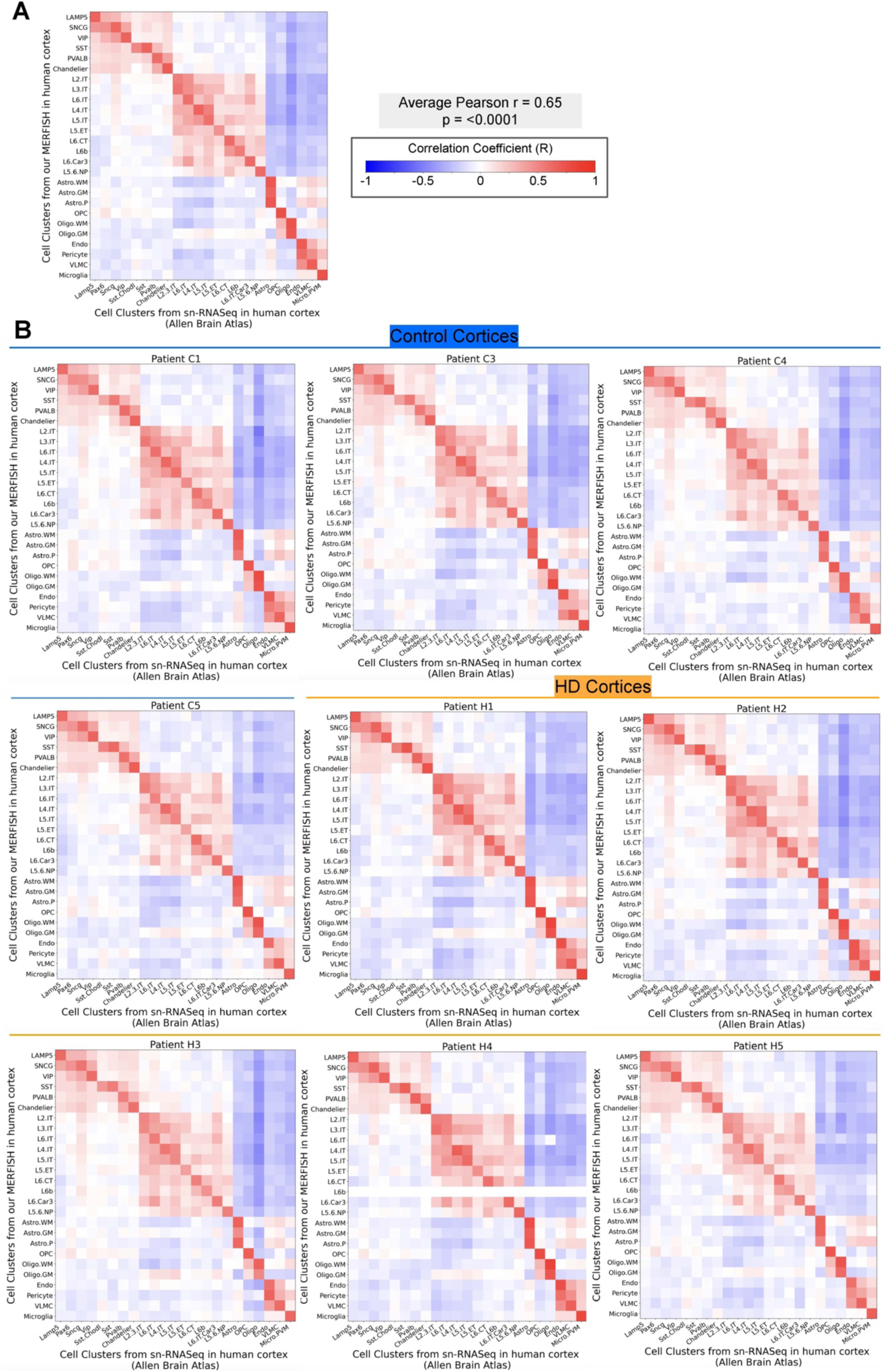
MERFISH clustering of 1128 gene measurements aligns with snRNA-seq. (**A**) Heatmap Pearson correlation plots between the major cell type clusters in our MERFISH clustering results in human cortex and snRNA-seq performed by the Allen Brain Atlas. (**A**) Heatmap Pearson correlation plots between the major cell type clusters in our MERFISH+ results in each cortical sample and snRNA-seq performed by the Allen Brain Atlas.

**Supplementary Figure S4:**
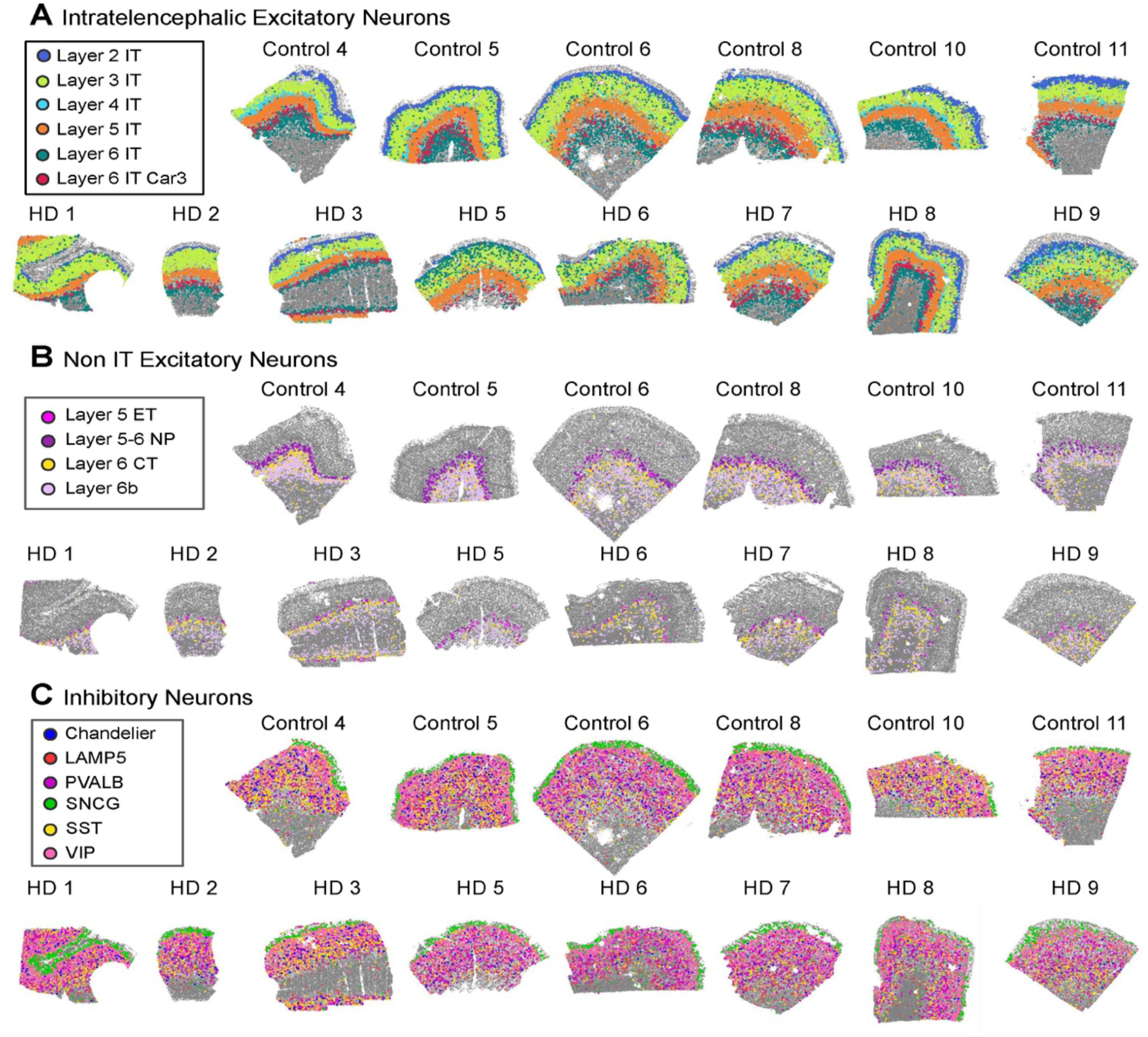
MERFISH spatial map of neuronal subtypes in HD and Control cortices. (**A**) Intratelencephalic Excitatory Neurons (Layer 2, 3, 4, 5, 6 IT) and Layer 6 IT Car3+ (Layer 6 Car3). (**B**) Non IT Excitatory Neurons: Layer 5 Extratelencephalic (Layer 5 ET), Layer 5/6 Near Projecting (Layer 5-6 NP), Layer 6 Corticothalamic (Layer 6 CT) and Layer 6b. (**C**) Inhibitory Neurons.

**Supplementary Figure S5:**
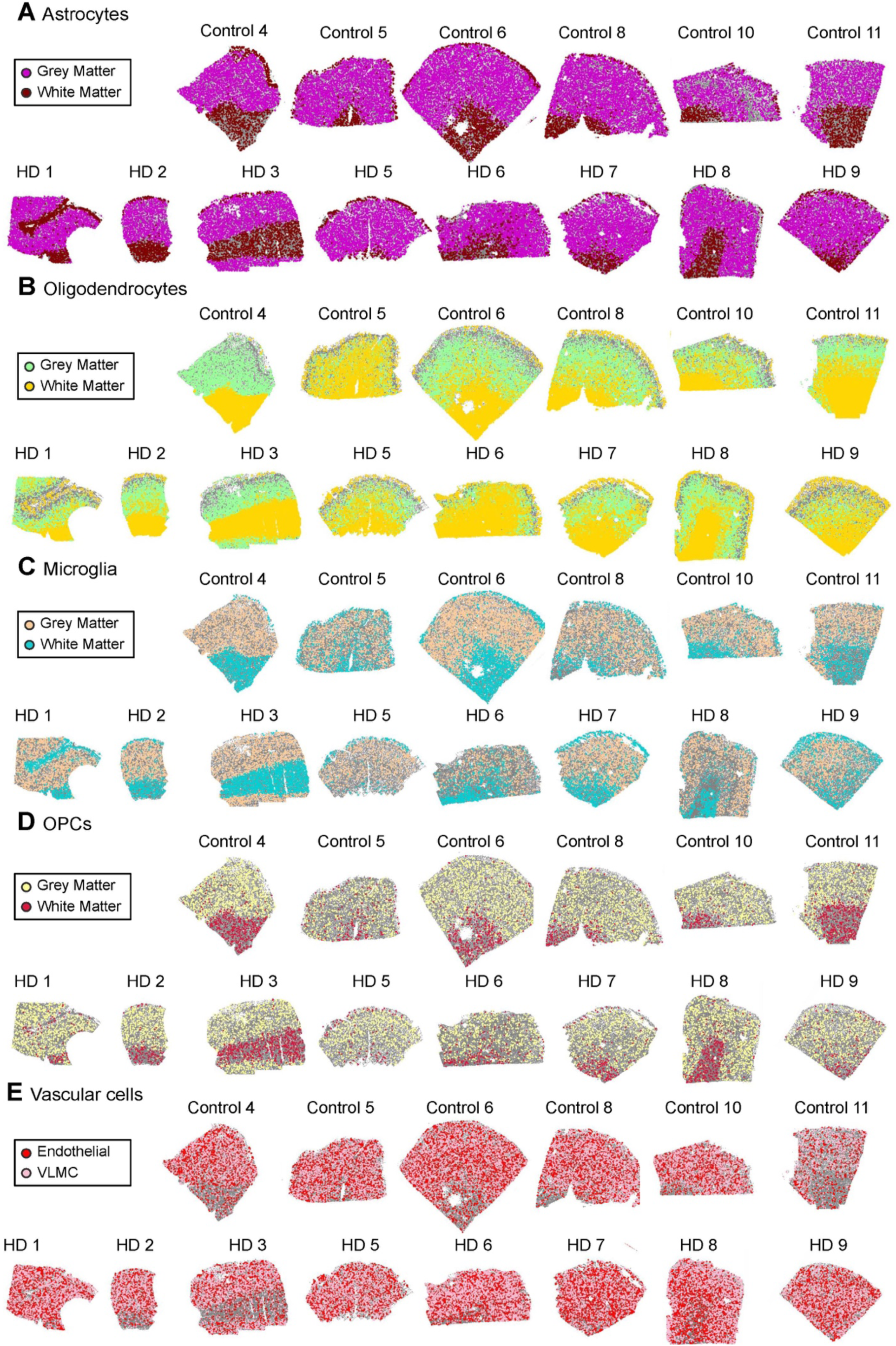
MERFISH spatial map of non-neuronal subtypes in HD and Control cortices. (**A**) Astrocytes. (**B**) Oligodendrocytes. (**C**) Microglia. (**D**) OPCs (oligodendrocytes precursor cells). (**E**) Vascular cells.

**Supplementary Figure S6:**
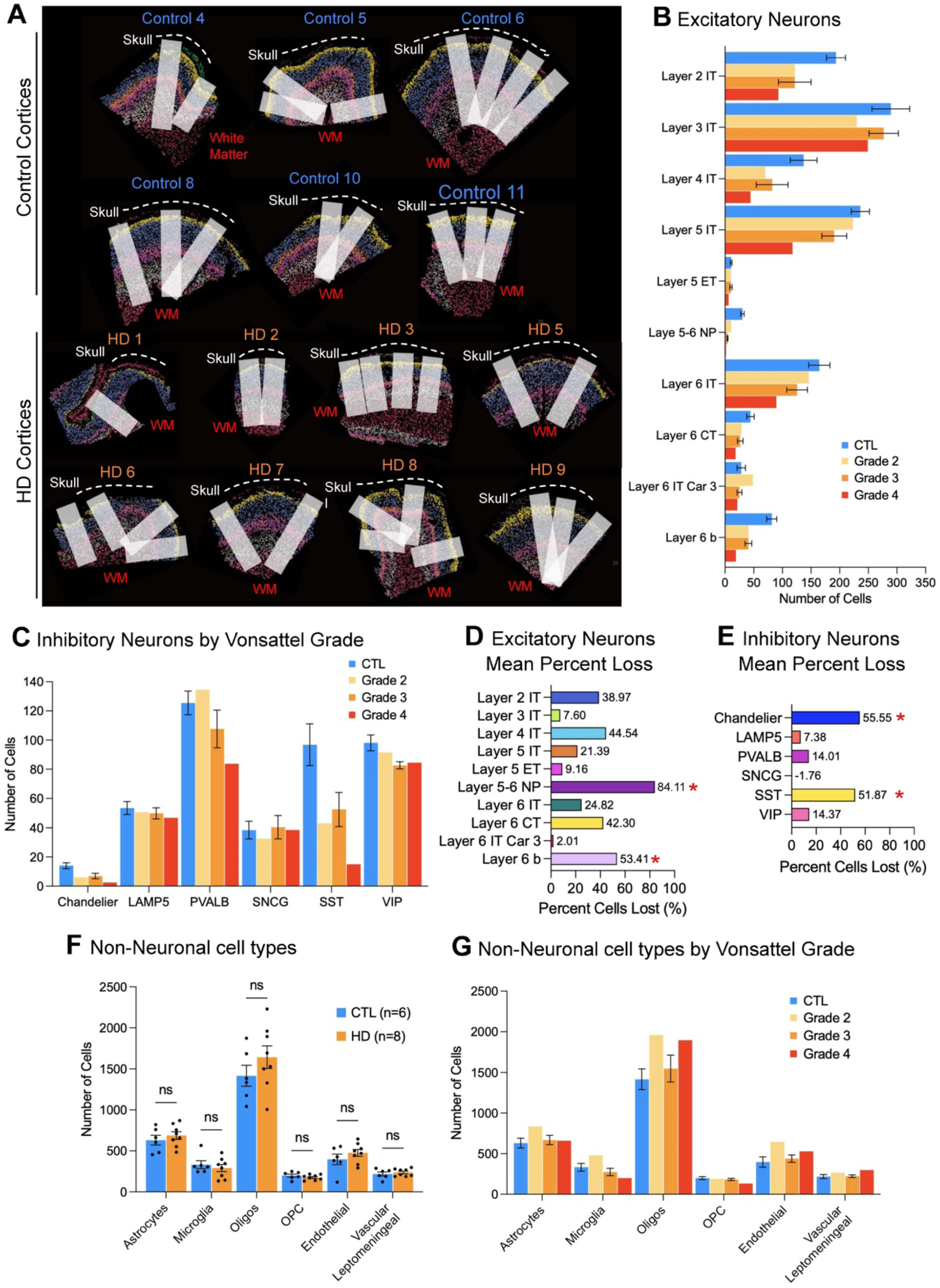
Cell type abundance in HD and Control cortex. (**A**) Selected columns within quantifications were performed across samples. (**B**) Quantification of Excitatory Neurons by Vonsattel Grade. (**C**) Quantification of Inhibitory Neurons by Vonsattel Grade. (**D**) Quantification of percentage of excitatory neurons lost. (**E**) Quantification of percentage of inhibitory neurons lost. (**F**) Quantification of non-neuronal cell types. (**G**) Quantification of non-neuronal cell types by Vonsattel Grade.

**Supplementary Figure S7:**
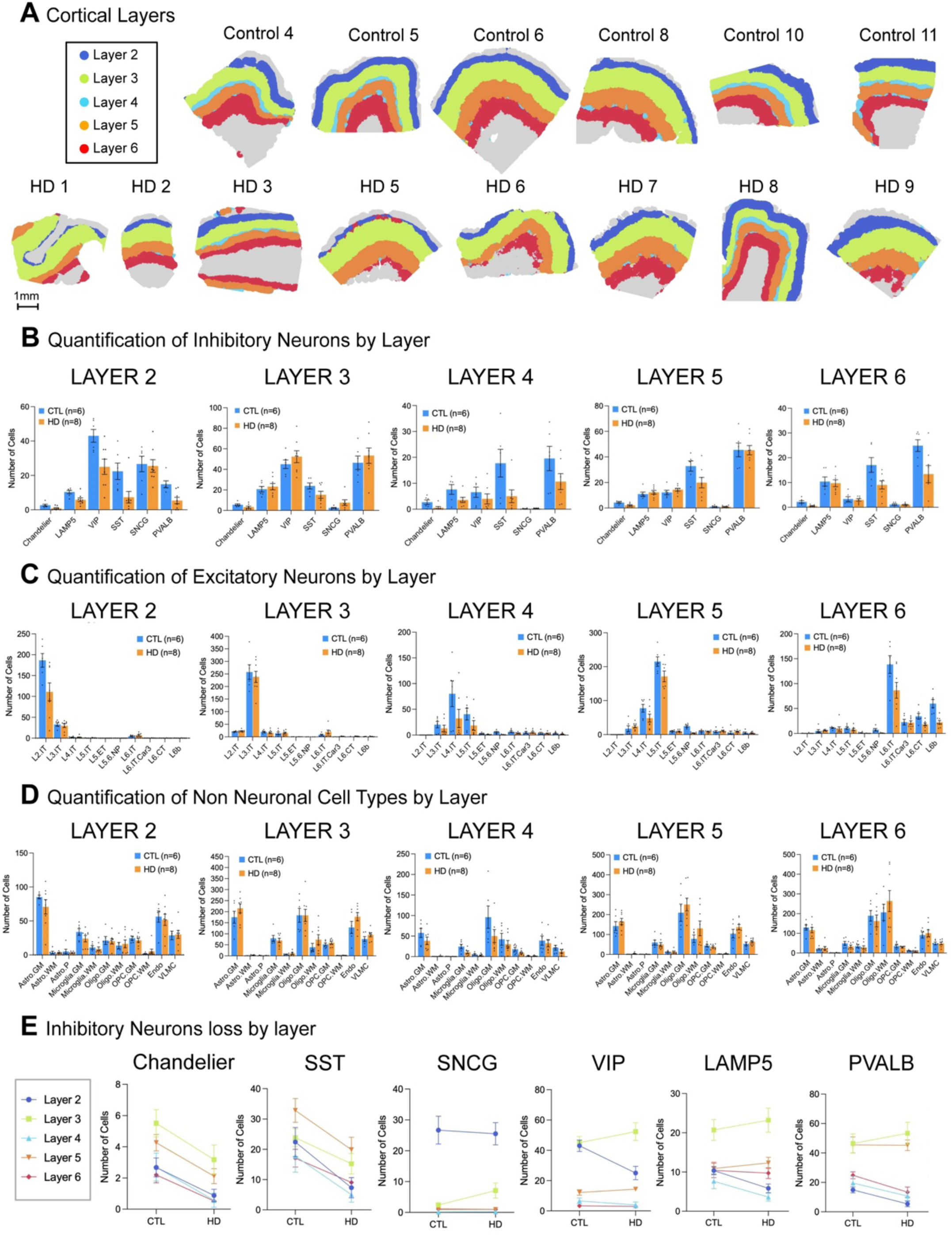
Cell type composition of each cortical layer. (**A**) Spatial map of non-neurological control and HD frontal cortices crossection showing the area occupied by each cortical layers as defined by their excitatory neuron composition. (**B**) Quantification of inhibitory neurons in each cortical layer. (**C**) Quantification of excitatory neurons in each cortical layer. (**D**) Quantification of non-neuronal cell types in each cortical layer.

**Supplementary Figure S8:**
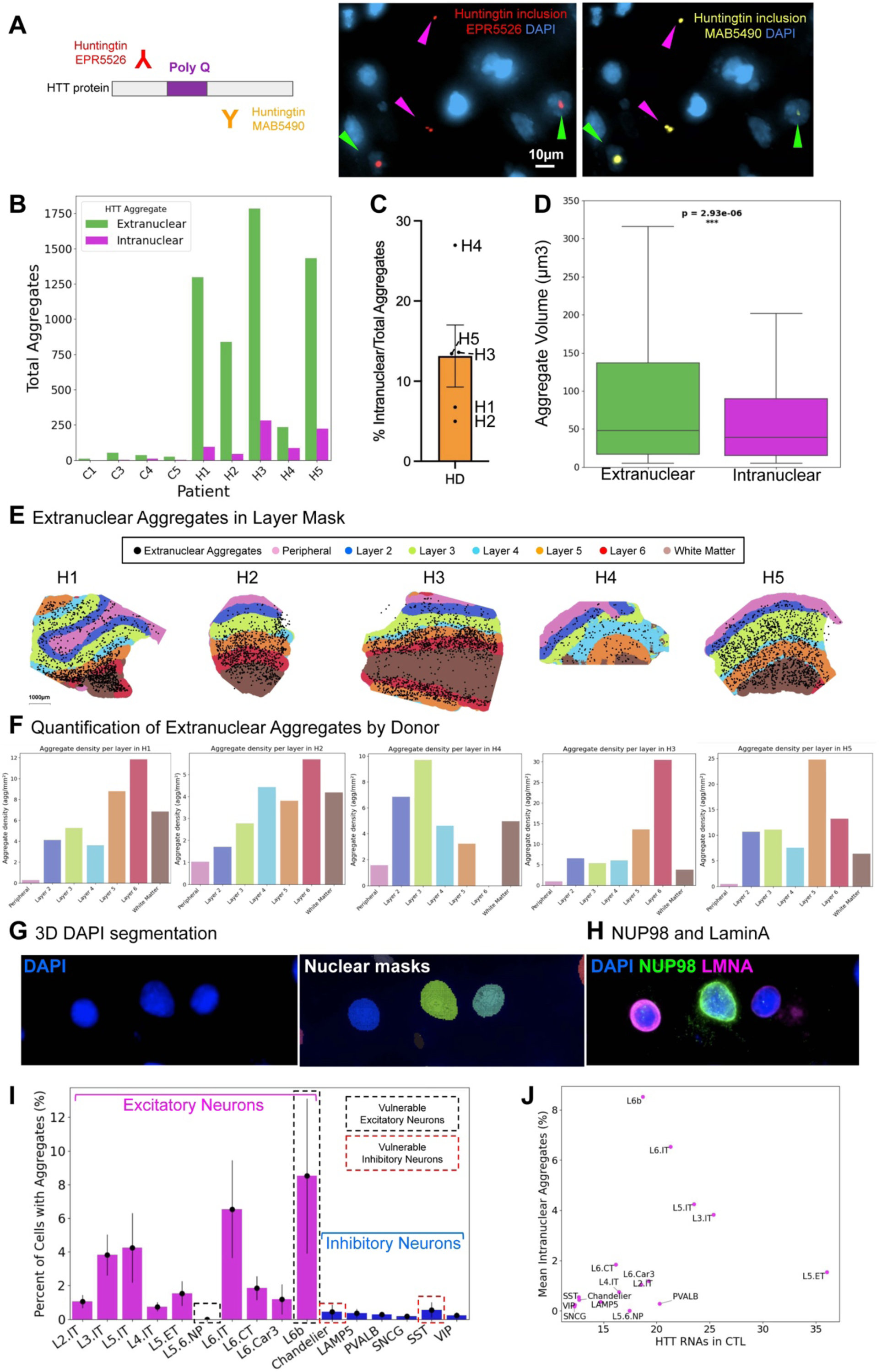
Characterization of intra- and extranuclear huntingtin aggregates. (**A**) Cartoon depicting the antibodies use to detect huntingtin and micrographs using them sequentially. The epitope of the antibody we used for downstream analysis (amino acids 115–129 of the huntingtin (HTT) protein) lies near the N-terminus just downstream of the polyglutamine (polyQ) and polyproline (polyP) tracts encoded by the CAG and CCG repeats, respectively. (**B**) Quantification of number of intra-(magenta) and extranuclear (green) HTT aggregates in HD and Control cortices. (**C**) Percentage of intranuclear HTT aggregates in HD cortices. (**D**) Intra-and extranuclear HTT aggregate volume (μm^3^) in HD cortices. (**I**) Spatial map of molecular define cell types from Layer 1 (peripheral), Layer, 2-6 and white matter in HD cortices and identified extranuclear aggregates with cortical layers masks. (**J**) Quantification of extranuclear aggregate density (aggregate per mm^2^) in each cortical layer mask in each individual HD cortex. (**G**) Example of nuclear 3D segmentation using DAPI. (**H**) Example of nuclear 3D segmentation using NUP98, Lamin A and DAPI. (**I**) Percentage of cells with intranuclear HTT aggregates in the neuronal clusters. (**J**) Correlation between the percentage of cells with intranuclear aggregates and the percentage cell loss in the neuronal clusters.

**Supplementary Figure S9:**
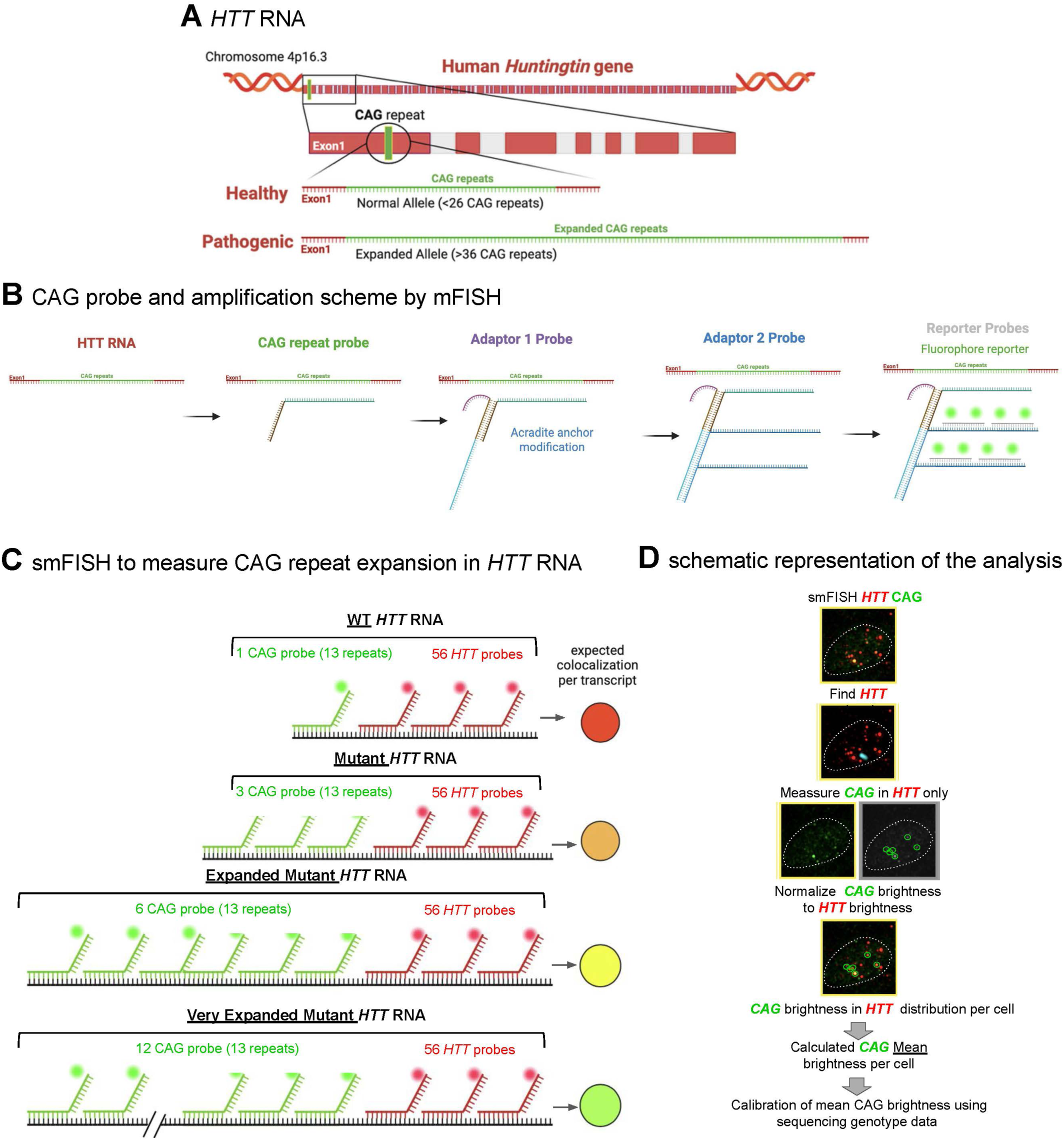
smFISH approach to quantify CAG repeats in *HTT* transcripts. (**A**) Illustration of *HTT* gene. (**B**) Illustration of the branching tree amplification approach for the CAG probe. (**C**) Illustration of the expected colocalization between CAG and *HTT* smFISH probes upon increasing amount of CAG repeats in *HTT* RNA. (**D**) Schematic representation of the analyses pipeline for CAG and *HTT* smFISH.

**Supplementary Figure S10:**
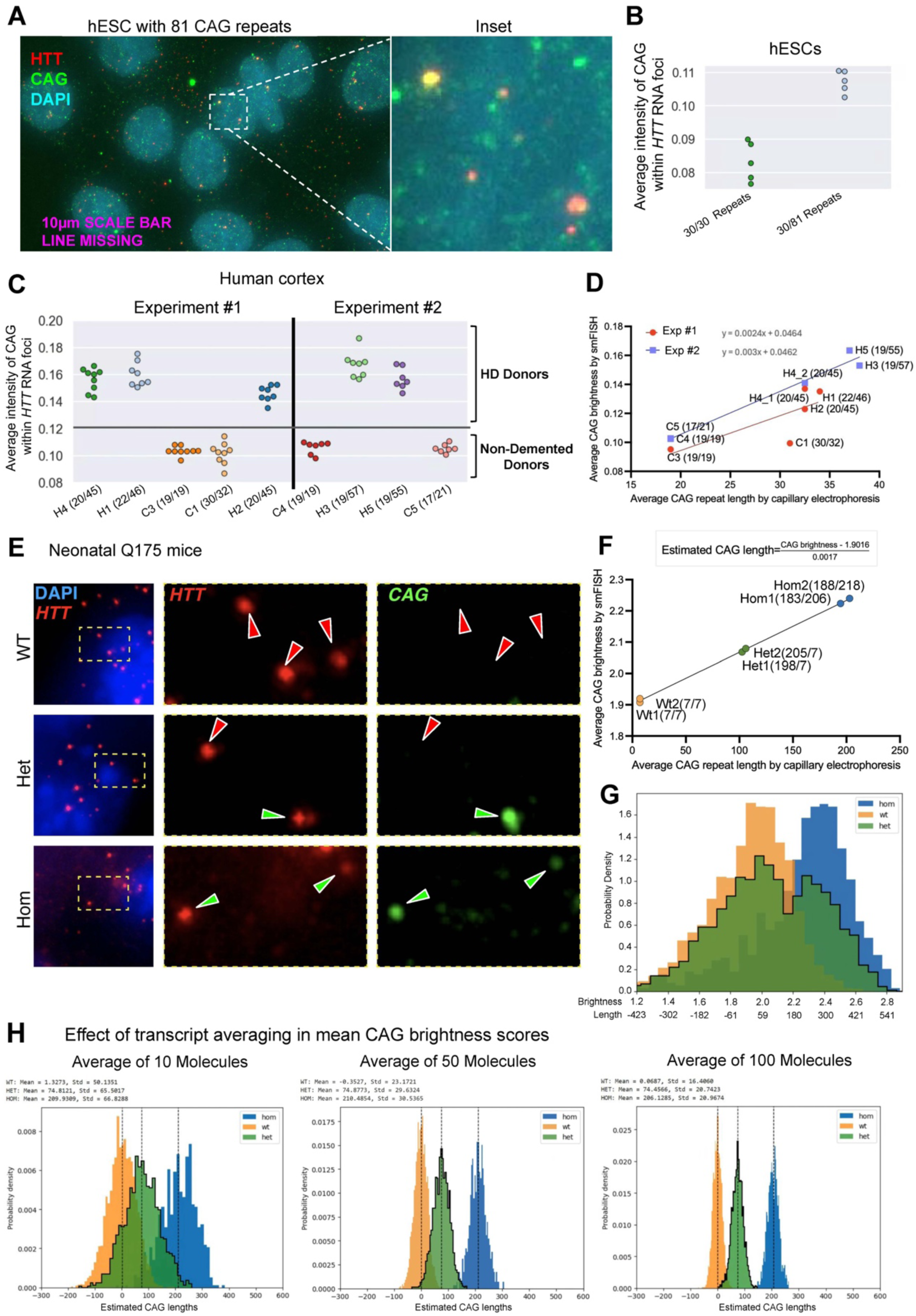
Calibrating the smFISH measurements of CAG repeats in *HTT* transcripts. (**A**) Representative image of *HTT* transcripts (red) and CAG probe (green) in human embryonic stem cells (hESC) with 30 and 81 CAG repeats in each *HTT* allele. DAPI in blue. (**B**) Quantification of the average intensity of CAG within *HTT* transcripts in each field of view of hESCs with 30/30 repeats and 30/81 repeats in HTT. (**C**) Quantification of the average intensity of CAG within *HTT* transcripts in each field of view of HD and control cortices. (**D**) Correlation between the calibrated average CAG repeats in *HTT* transcripts measured by smFISH with the average CAG repeat length measured by capillarity electrophoresis in control and HD cortices. (**E**) Representative images of *HTT* transcripts (red) and CAG probe (green) in WT, Het and Homo Htt Q175 neonatal mice. DAPI in blue. (**F**) Correlation between the average CAG brightness in *HTT* transcripts by smFISH and the average CAG repeat length measured by capillarity electrophoresis. (**G**) Probability density of CAG brightness in each in *HTT* transcript by smFISH. (**H**) Probability density of CAG brightness when averaging 5, 10 or 100 *HTT* transcripts by smFISH.

**Supplementary Figure S11:**
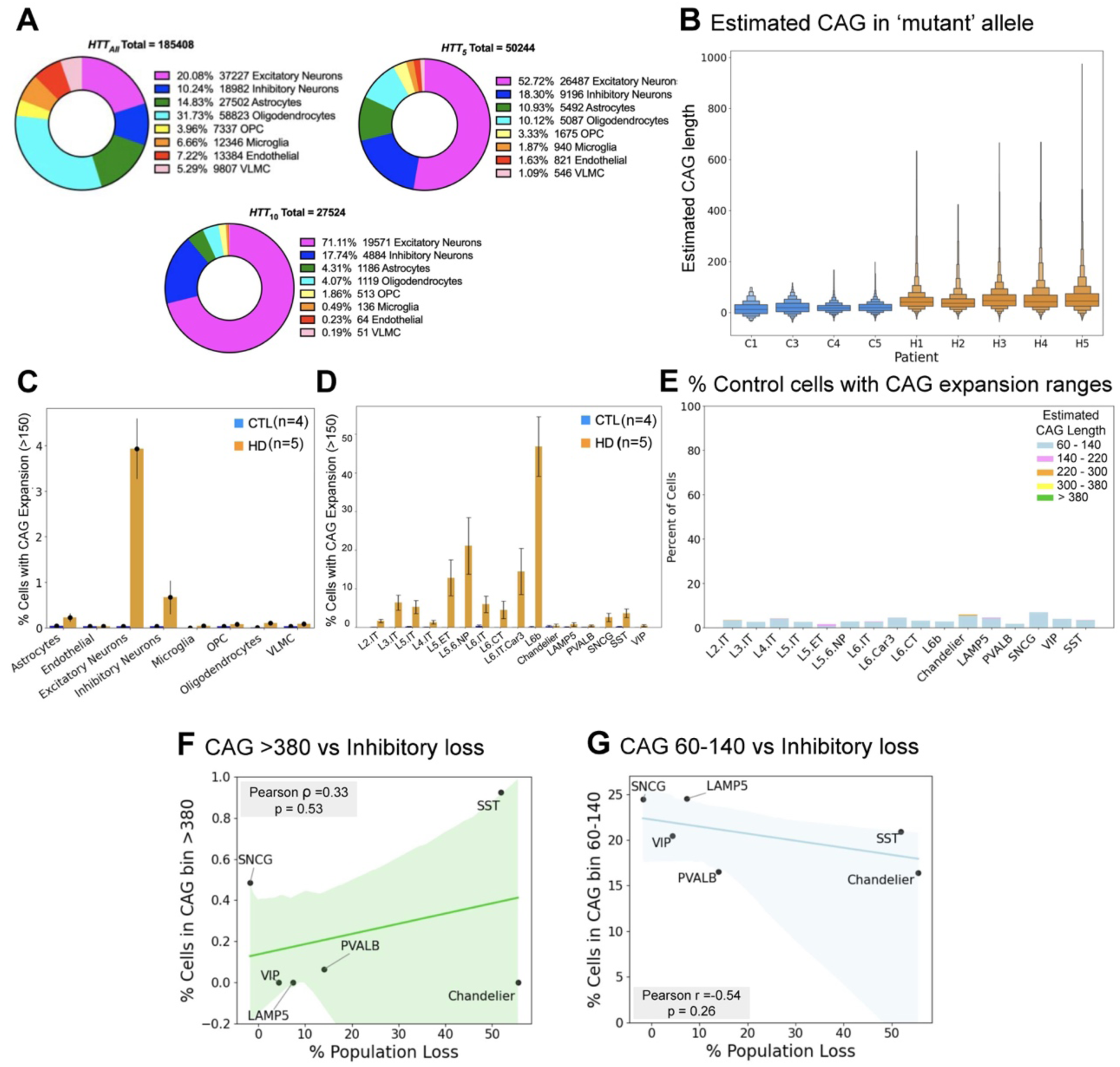
Characterization of CAG repeats in HTT transcripts in HD and control cortex. (A) Percentage of the different cell populations in human cortex (*HTT_All_*), percentage of the different cell populations when filtering of cell that contain at least 5 HTT transcripts (*HTT_5_*) and 10 HTT transcripts (*HTT_10_*). (B) Estimated CAG repeats in *HTT* RNA per cell in control and HD cortices. (**C**) Quantification of percentage of cells in the main clusters that have an estimated CAG length larger than 150 repeats in 5 HD and 4 control frontal cortices. (**D**) Quantification of percentage of cells in the neuronal clusters that have an estimated CAG length larger than 150 repeats in 5 HD and 4 control frontal cortices. (**E**) Quantification of percentage of cells in the neuronal clusters that have 60-140, 140-220, 220-300,300-380 or more than 380 estimated CAG repeats in *HTT* in 4 control frontal cortices. (**F-G**) Correlation between the percentage of cell in each inhibitory neuronal subtype that have an estimated CAG length of >380 (F) and 60-140 (G) repeats and the percentage of population loss.

**Supplementary Figure S12:**
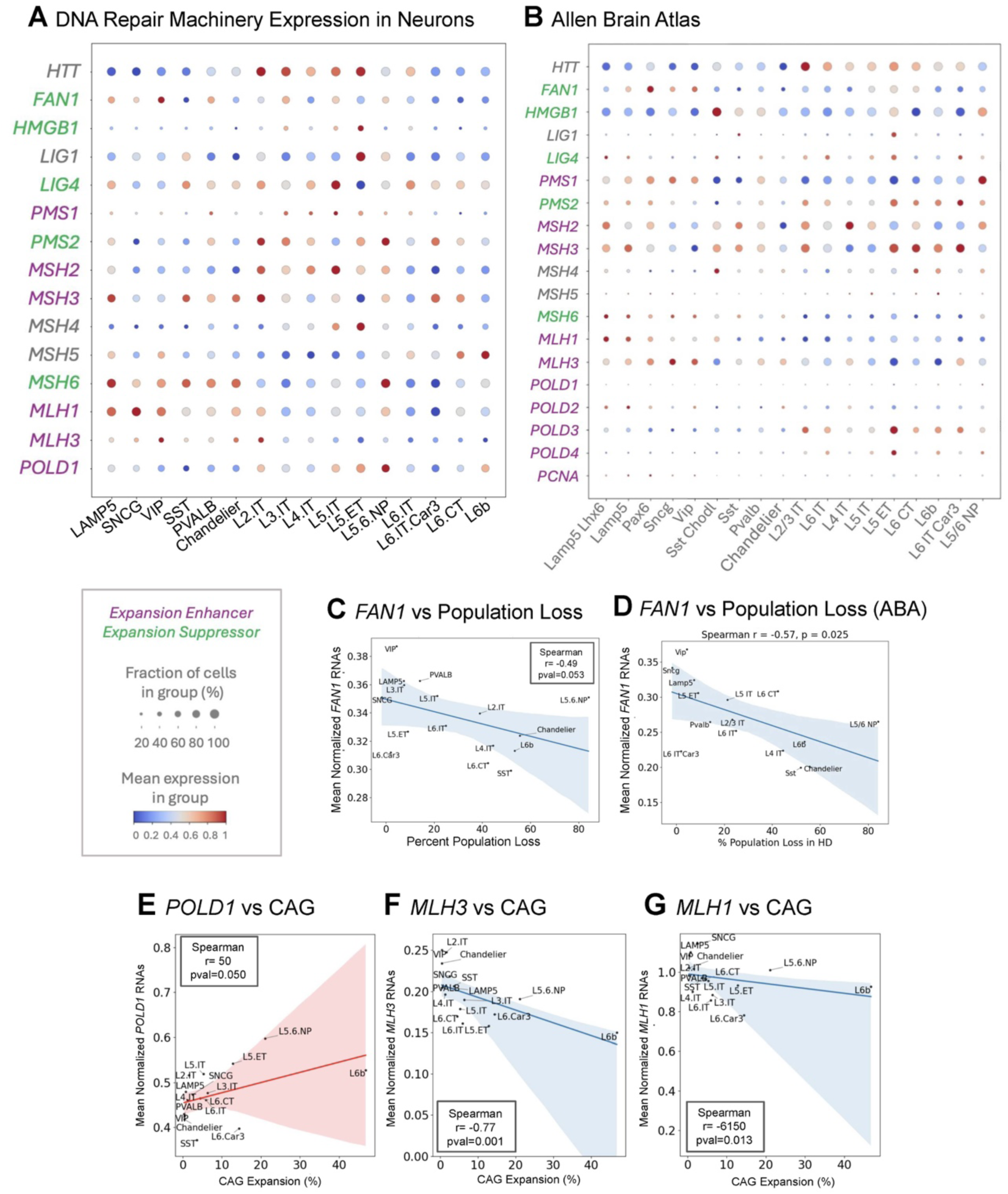
Correlation between cell type vulnerability, somatic repeat expansion and expression of HD related DNA repair genes. (**A**) Dot plot of DNA repair genes involved in HD expressed in neuronal clusters from our MERFISH data in human control cortex. (**B**) Dot plot of DNA repair genes involved in HD expressed in neuronal clusters from single-cell sequencing cell atlases of the human cortex^25^. Magenta genes are known to be CAG expansion enhancer and green genes expansion suppressor^12^. (**C**) Correlation for *FAN1* expression levels and percentage lost measured in our Multimodal MERFISH. (**D**) Correlation for *FAN1* expression levels measured by single-cell sequencing in human cortex^25^ and percentage lost measured in our Multimodal MERFISH. (**E**) Correlation for *POLD1* expression levels and percentage of cells that have an estimated CAG length larger than 150 repeats in Multimodal MERFISH. (**F**) Correlation for *MLH3* expression levels and percentage of cells that have an estimated CAG length larger than 150 repeats in Multimodal MERFISH. (**G**) Correlation for *MLH1* expression levels and percentage of cells that have an estimated CAG length larger than 150 repeats in Multimodal MERFISH.

**Supplementary Figure S13:**
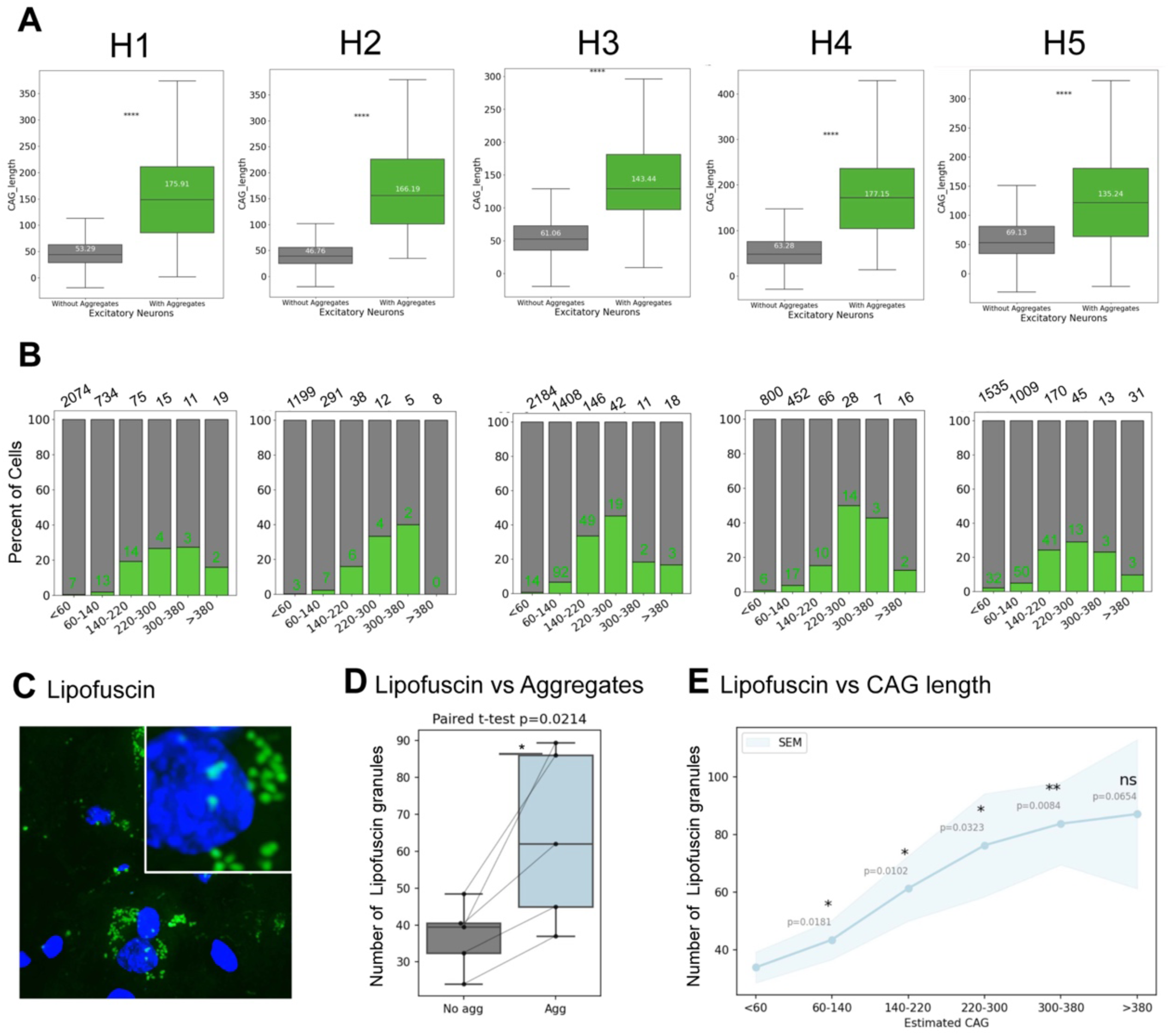
Intranuclear aggregation and lipofuscin granule number increase with somatic repeat expansion. (**A**) Estimated CAG length in *HTT* RNA in cells without or with intranuclear HTT aggregates in each HD cortex. (**B**) Percentage of cells with and without intranuclear HTT aggregates across different estimated CAG length in *HTT* RNA per cell in each HD cortex. **(C)** Representative image of lipofuscin in HD cortex. **(D)** Quantification of lipofuscin granules in excitatory neurons without or with intranuclear aggregation. **(E)** Quantification of lipofuscin granules in excitatory neurons that have 60-140, 140-220, 220-300,300-380 or more than 380 estimated CAG repeats in *HTT* in HD cortices.

**Supplementary Figure S14:**
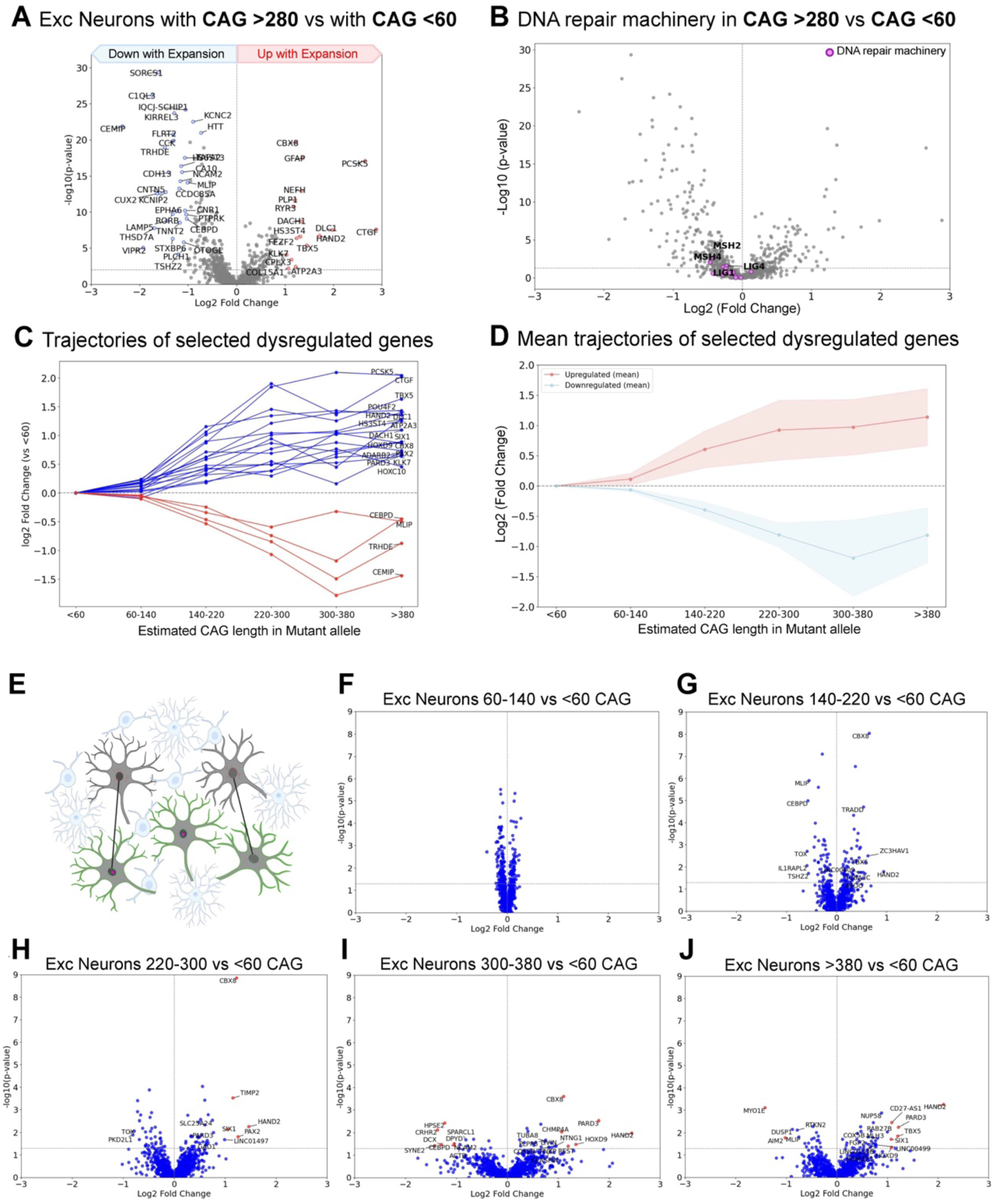
Transcriptional changes associated with large somatic repeat expansions. (**A**) Volcano plot of statistical significance versus the log2 fold change for gene expression in neurons with large somatic repeat expansion (estimated CAG length >280 repeats) compared to their nearest neighbors of the same cell type that lacked large expansion (estimated CAG length <60 repeats). (**B**) Volcano in (A) with genes involved in DNA damage and repair labelled. (**C**) Plot of the log2 fold change for genes differentially expressed in neurons with increasing extend of somatic repeat expansion (<60, 60-140, 140-220, 220-300,300-380 or more than 380 estimated CAG repeats in *HTT)*. (**D**) Plot of the mean log2 fold change for genes differentially expressed (in C) in neurons with increasing extend of somatic repeat expansion (<60, 60-140, 140-220, 220-300,300-380 or more than 380 estimated CAG repeats in *HTT)*. (**E**) Schematic cartoon of the closest neighbor analysis used to a series of pairwise comparisons between individual cells. **(F-J)** Volcano plot of statistical significance versus the log2 fold change for gene expression in neurons with large somatic expansions [60-140 (F),140-220 (G), 220-300 (H), 300-380 (I) and >380 (J) estimated CAG repeats] compared to their nearest neighbors of the same cell type that lacked large somatic expansion (less than 60 estimated CAG repeats in *HTT)*.

## Supplemental Table Legends

**Table S1. Sequences of the Multimodal MERFISH probes and associated primers/readouts**

**Table S2. Metadata for brain donors whose data contributed to this study**

**Table S3. Mean *HTT* transcripts and CAG error per cell measurements per each neuronal cluster**

